## Supplementary Methods for "Histone modifications during the life cycle of the brown alga *Ectocarpus*"

**Liquid chromatography-MS/MS Analysis**

After precipitation, histone preparations were separated on 14% acrylamide SDS–PAGE gels. Excised histone bands were washed and the proteins reduced with 10 mM DTT prior to alkylation with 55 mM chloroacetamide. After washing and shrinking of the gel fragments with 100% acetonitrile, in-gel digestion was carried out using trypsin/LysC (0.1µg) overnight in 25 mM ammonium bicarbonate at 30°C. Peptides were then extracted using 60/35/5 MeCN/H2O/HCOOH, vacuum concentrated to dryness and reconstituted in loading buffer A (2/98 MeCN/H2O + 0.05% TFA) prior to liquid chromatography-tandem mass spectrometry (LC-MS/MS) analysis.

The extracted peptides were separated chromatographically using an RSLCnano system (Ultimate 3000, Thermo Scientific) coupled with a TripleTOF™ 6600 mass spectrometer (ABSciex) or a Q Exactive HF-X mass spectrometer (Thermo Scientific).

For the tripleTOF™ 6600 analysis, peptides were loaded onto a C18-reversed phase column (300-µm inner diameter × 5 mm; C18 PepMap™ 100Å pore size, 5µm granulometry, Thermo Scientific) with loading buffer A, separated and MS data acquired using the Analyst software. Peptide separation was carried out on a 50 cm x 75-µm C18 column (nanoViper Acclaim PepMap™ RSLC, 3 µm, 100Å, Thermo Scientific) over a linear gradient of 137 min from 1% to 20% or 1% to 35% of buffer B admixed to buffer A' (buffer B: 100% MeCN and buffer A': 2/98 MeCN/H2O + 0.1% HCOOH) at a flow rate of 300 nL/min. The mass spectrometer was operated in DDA top20 or top40 high sensitivity mode with 250 and 100 or 120 ms acquisition time for MS1 and MS2 scans, respectively, and 10 s or 30 s dynamic exclusion, with or without an inclusion list.

For the Q Exactive HF-X analysis, peptides were loaded onto a C18-reversed phase column (75-µm inner diameter × 2 cm; nanoViper Acclaim PepMap™ 100, Thermo Scientific) with buffer A', separated and MS data acquired using Xcalibur software. Peptide separation was carried out over a linear gradient of 91 min from 2% to 30% buffer B (75-µm inner diameter × 50 cm; nanoViper Acclaim PepMap™ RSLC, 2 µm, 100Å, Thermo Scientific) at a flow rate of 300 nL/min. Full-scan MS was acquired in the Orbitrap analyzer with a resolution set to 120,000 and the top20 intense ions were subjected to Orbitrap for futher fragmentation via high energy collision dissociation (HCD) activation and a dynamic exclusion on 20 s.

For identification, the data were queried against the UniProtKB/Swiss-Prot *Ectocarpus* database (19/10/2015, containing 17103 entries) using Mascot™ (version 2.5.1).

Carbamidomethylation of cysteines, oxidation of methionines, propionylation of lysines, acetylation of lysines and protein N-termini, methylation and dimethylation of lysines and arginines, trimethylation of lysines and ubiquitination of lysine were set as variable modifications and with a maximum of nine modifications for all Mascot searches. Specificity of trypsin digestion was set for cleavage after Lys or Arg except before Pro, and five missed trypsin cleavage sites were allowed. The mass tolerances in MS and MS/MS were set to 10 ppm and 0.02 ppm for the HF-X or 100 mmu for the 6600, respectively. The resulting Mascot files were further processed using myProMS v3.6 [1].The maximum false discovery rate (FDR) calculation was set to 2% at the peptide level for the whole study (QVALITY algorithm). The mass spectrometry proteomics data have been deposited with the ProteomeXchange Consortium via the PRIDE partner repository [2] with the dataset identifier PXD013535.

### References

1. Poullet P, Carpentier S, Barillot E. myProMS, a web server for management and validation of mass spectrometry-based proteomic data. Proteomics. 2007;7:2553–6.

2. Perez-Riverol Y, Csordas A, Bai J, Bernal-Llinares M, Hewapathirana S, Kundu DJ, et al. The PRIDE database and related tools and resources in 2019: improving support for quantification data. Nucleic Acids Res. 2019;47:D442–50.
