## Supplementary figures and images for "Histone modifications during the life cycle of the brown alga *Ectocarpus*"

### Fig S1 Histone clusters

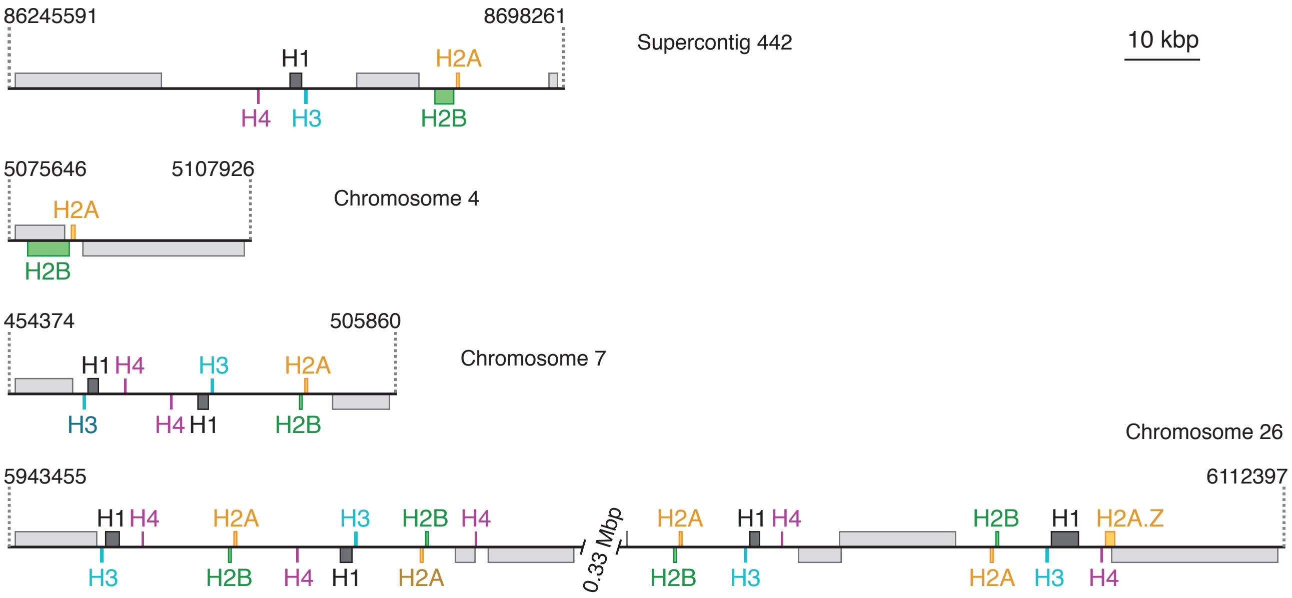
