## Supplementary material for "Histone modifications during the life cycle of the brown alga *Ectocarpus*": Fig S2 Mass Spectra

### Slide 1
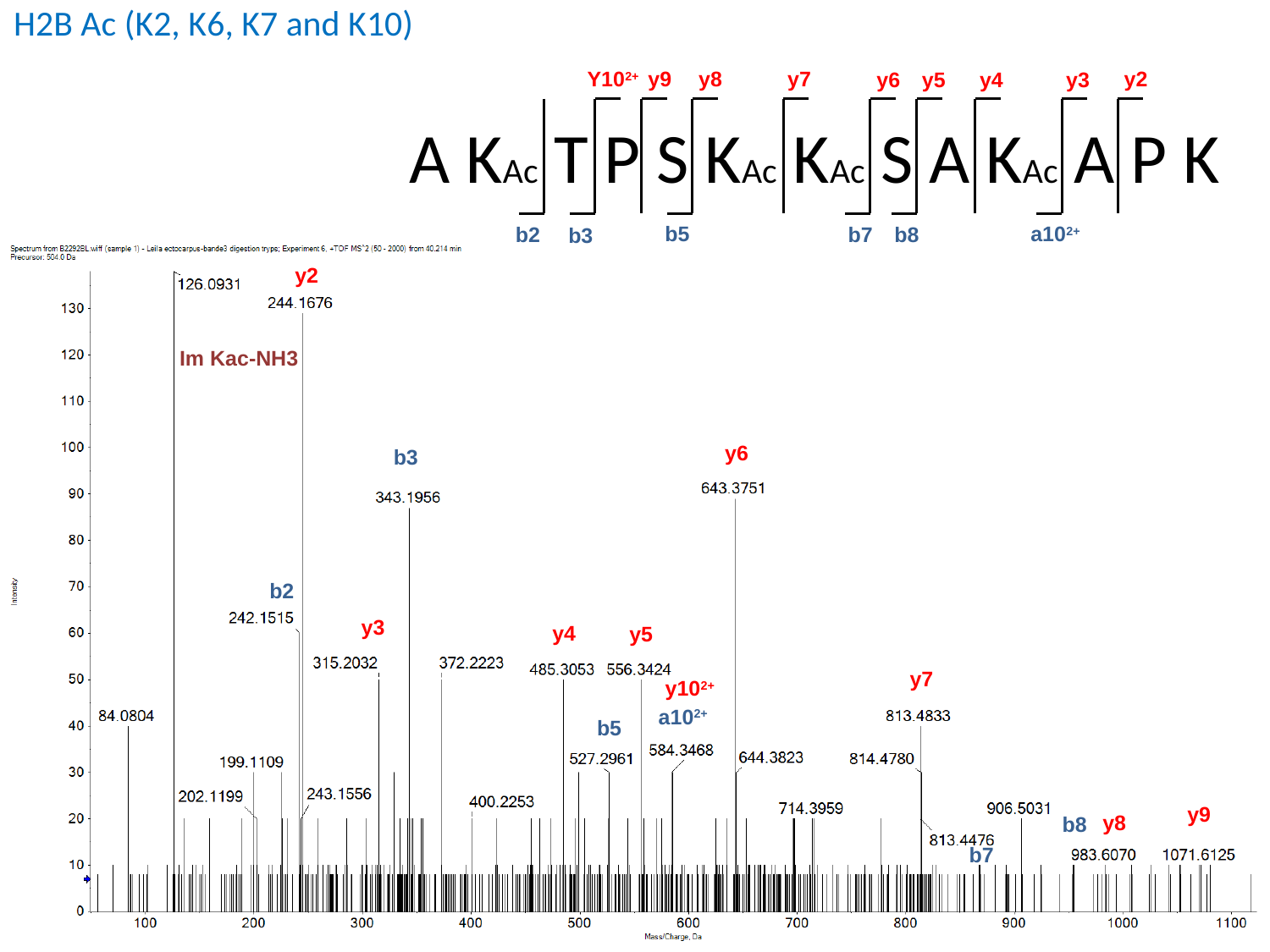

H2B Ac (K2, K6, K7 and K10)
y7
Y10^2+^
y8
y2
y9
y6
y5
y4
y3
A KAc T P S KAc KAc S A KAc A P K
a10^2+^
b5
b7
b2
b8
b3
y2
Im Kac-NH3
y6
b3
b2
y3
y4
y5
y7
a10^2+^
b5
y9
y8
b8
b7
y10^2+^

### Slide 2
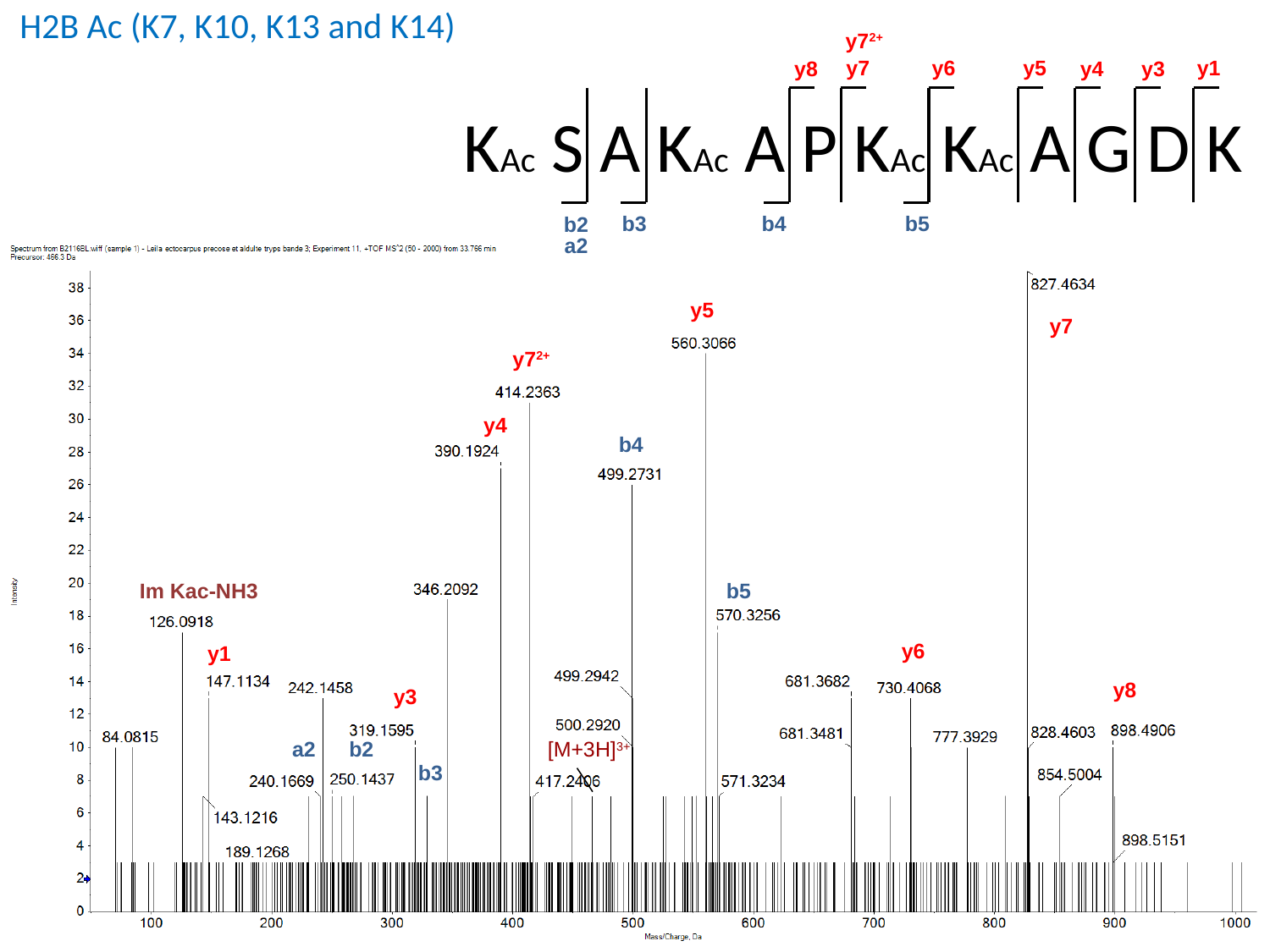

H2B Ac (K7, K10, K13 and K14)
y7^2+^
y7
y1
y5
y6
y8
y4
y3
KAc S A KAc A P KAc KAc A G D K
b4
b3
b5
b2
a2
y5
y7
y7^2+^
y4
b4
b5
Im Kac-NH3
y6
y1
y8
y3
[M+3H]^3+^
a2
b2
b3

### Slide 3
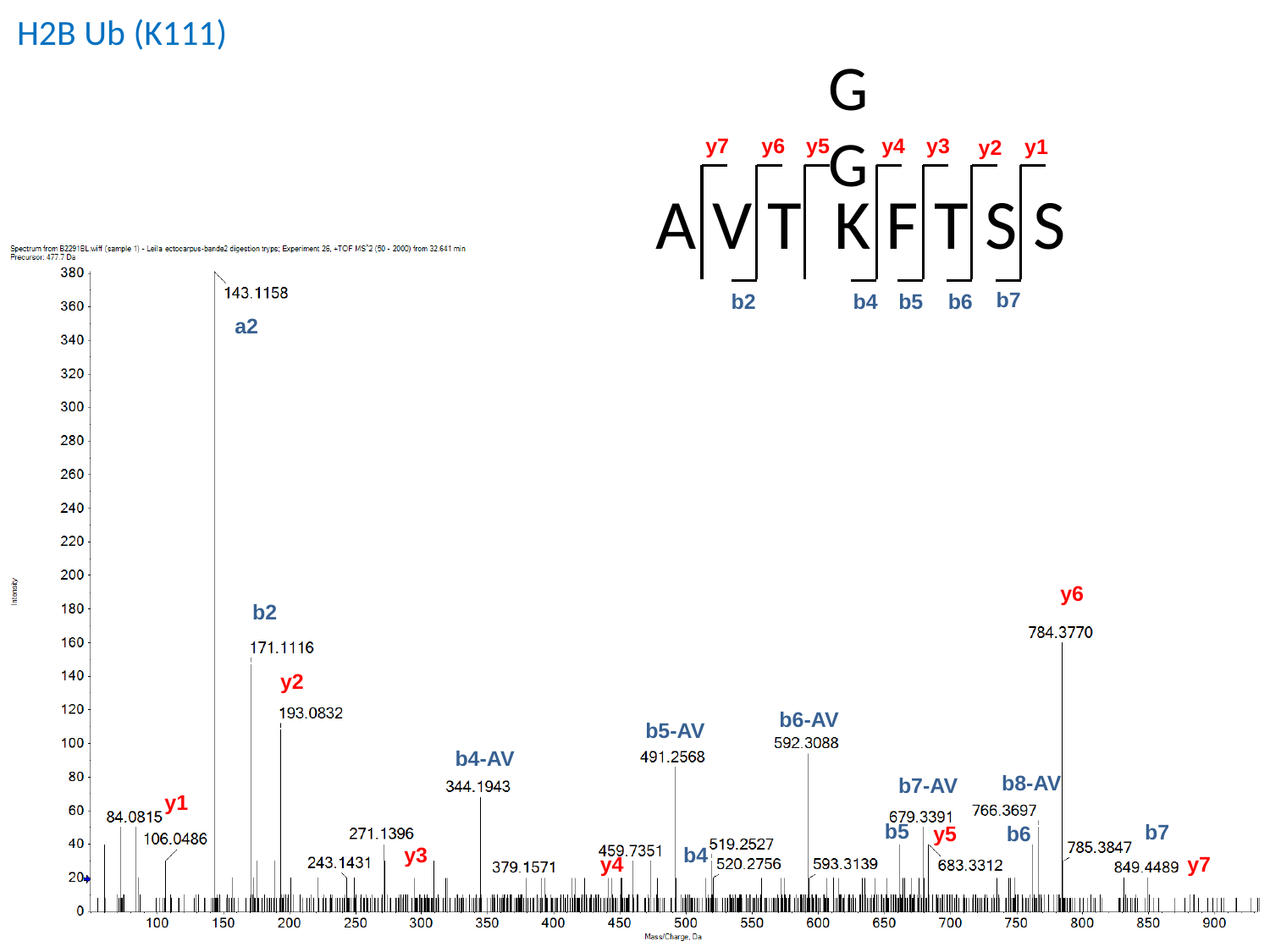

H2B Ub (K111)
GG
y7
y5
y4
y6
y3
y1
y2
A V T K F T S S
b7
b2
b4
b5
b6
a2
y6
b2
y2
b6-AV
b5-AV
b4-AV
b8-AV
b7-AV
y1
b5
b7
b6
y5
y3
b4
y4
y7

### Slide 4
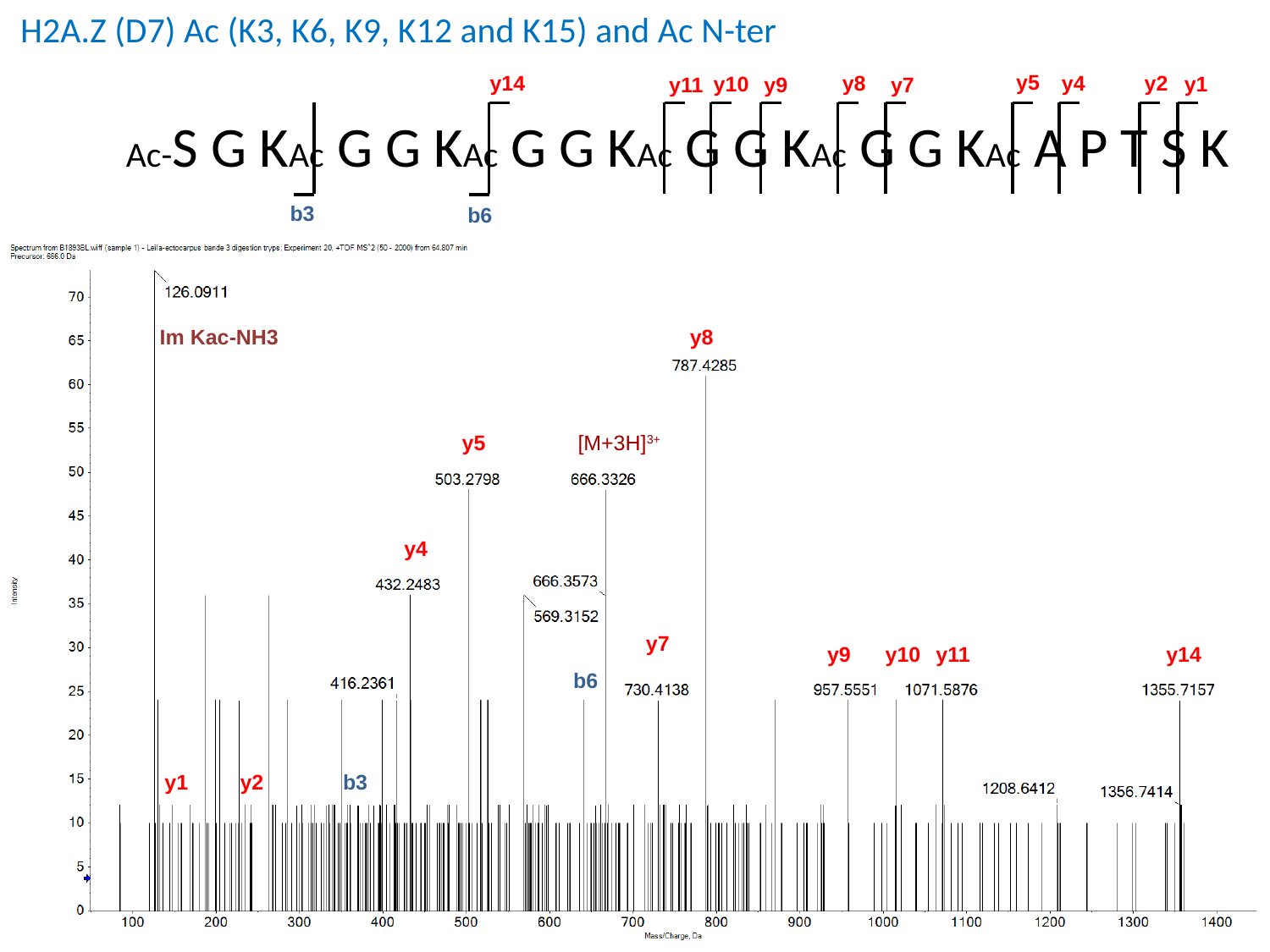

H2A.Z (D7) Ac (K3, K6, K9, K12 and K15) and Ac N-ter
y5
y14
y4
y8
y2
y1
y10
y9
y7
y11
b3
b6
Ac-S G KAc G G KAc G G KAc G G KAc G G KAc A P T S K
Im Kac-NH3
y8
[M+3H]^3+^
y5
y4
y7
y14
y11
y10
y9
b6
b3
y2
y1

### Slide 5
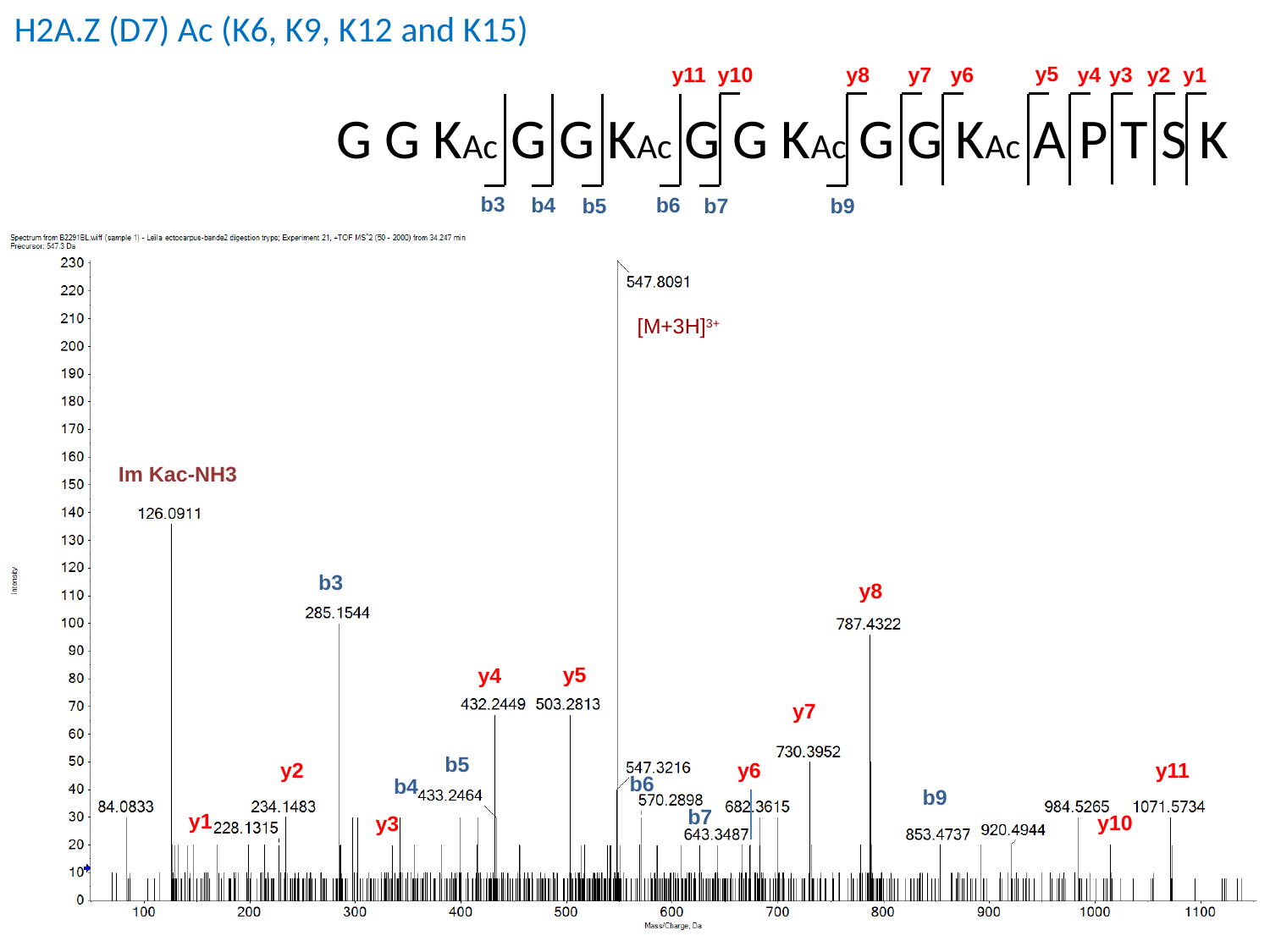

H2A.Z (D7) Ac (K6, K9, K12 and K15)
y5
y10
y4
y3
y6
y1
y11
y8
y7
y2
G G KAc G G KAc G G KAc G G KAc A P T S K
b3
b4
b6
b7
b5
b9
[M+3H]^3+^
Im Kac-NH3
b3
y8
y5
y4
y7
b5
y2
y6
y11
b6
b4
b9
b7
y1
y10
y3

### Slide 6
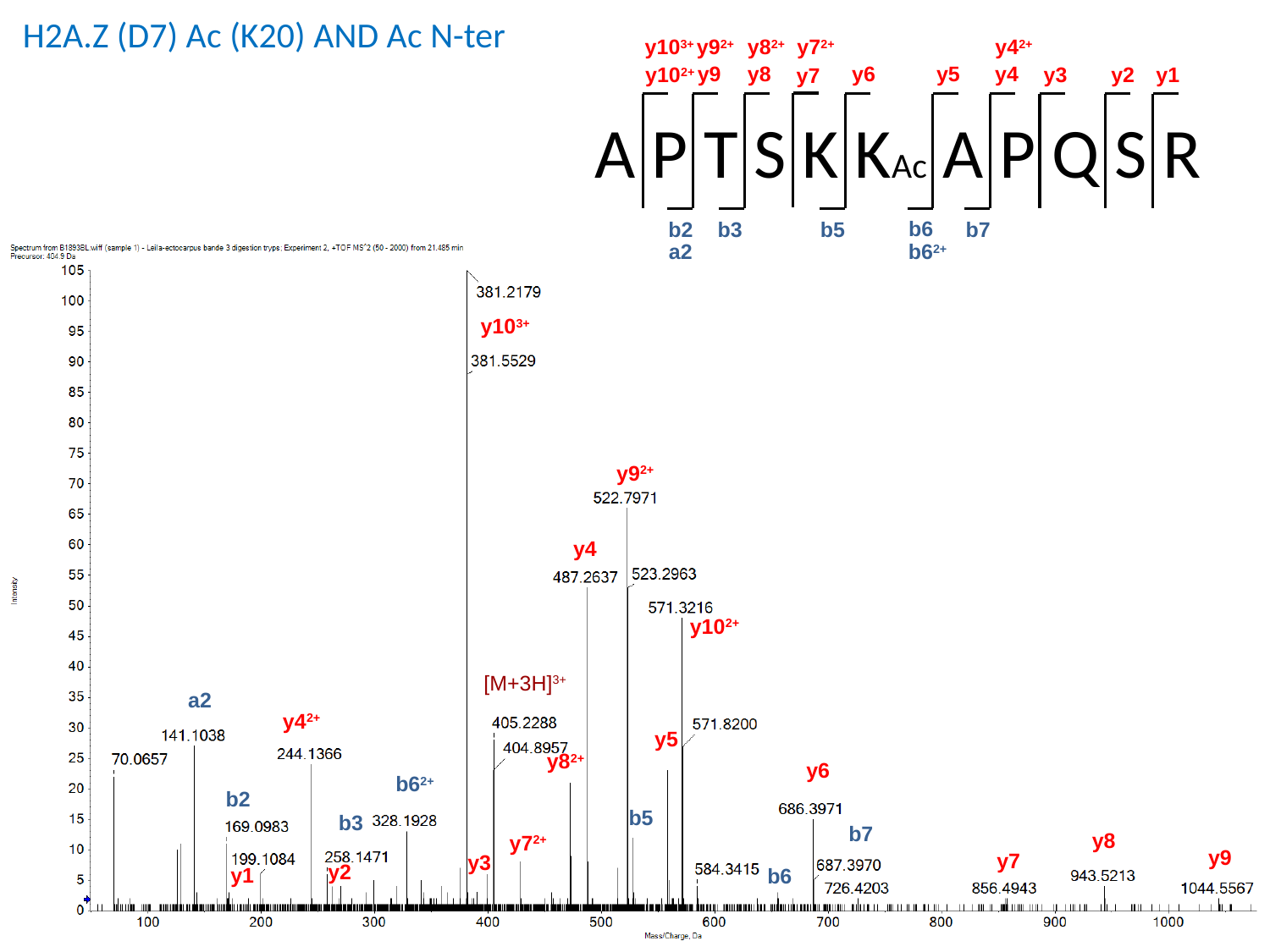

H2A.Z (D7) Ac (K20) AND Ac N-ter
y10^3+^
y9^2+^
y8^2+^
y7^2+^
y4^2+^
y8
y6
y9
y5
y4
y10^2+^
y3
y2
y1
y7
A P T S K KAc A P Q S R
b6
b3
b2
b7
b5
a2
b6^2+^
y10^3+^
y9^2+^
y4
y10^2+^
[M+3H]^3+^
a2
y4^2+^
y5
y8^2+^
y6
b6^2+^
b2
b5
b3
b7
y8
y7^2+^
y9
y7
y3
y2
y1
b6

### Slide 7
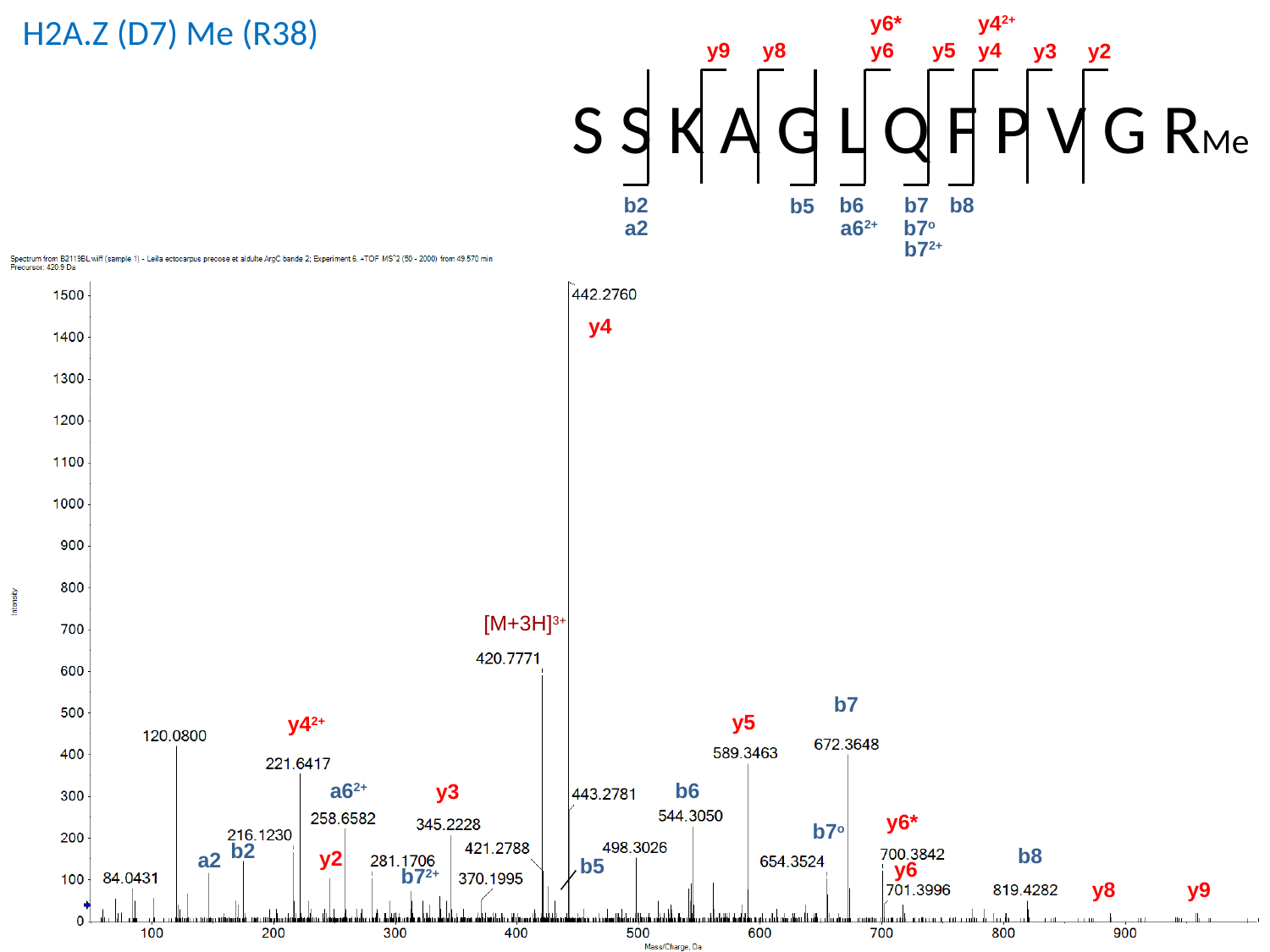

y6*
y4^2+^
y9
y8
y6
y5
y4
y3
y2
S S K A G L Q F P V G RMe
b8
b7
b6
b2
b5
a2
a6^2+^
b7^o^
b7^2+^
H2A.Z (D7) Me (R38)
y4
[M+3H]^3+^
b7
y5
y4^2+^
a6^2+^
b6
y3
y6*
b7^o^
b2
b8
y2
a2
b5
y6
b7^2+^
y9
y8

### Slide 8
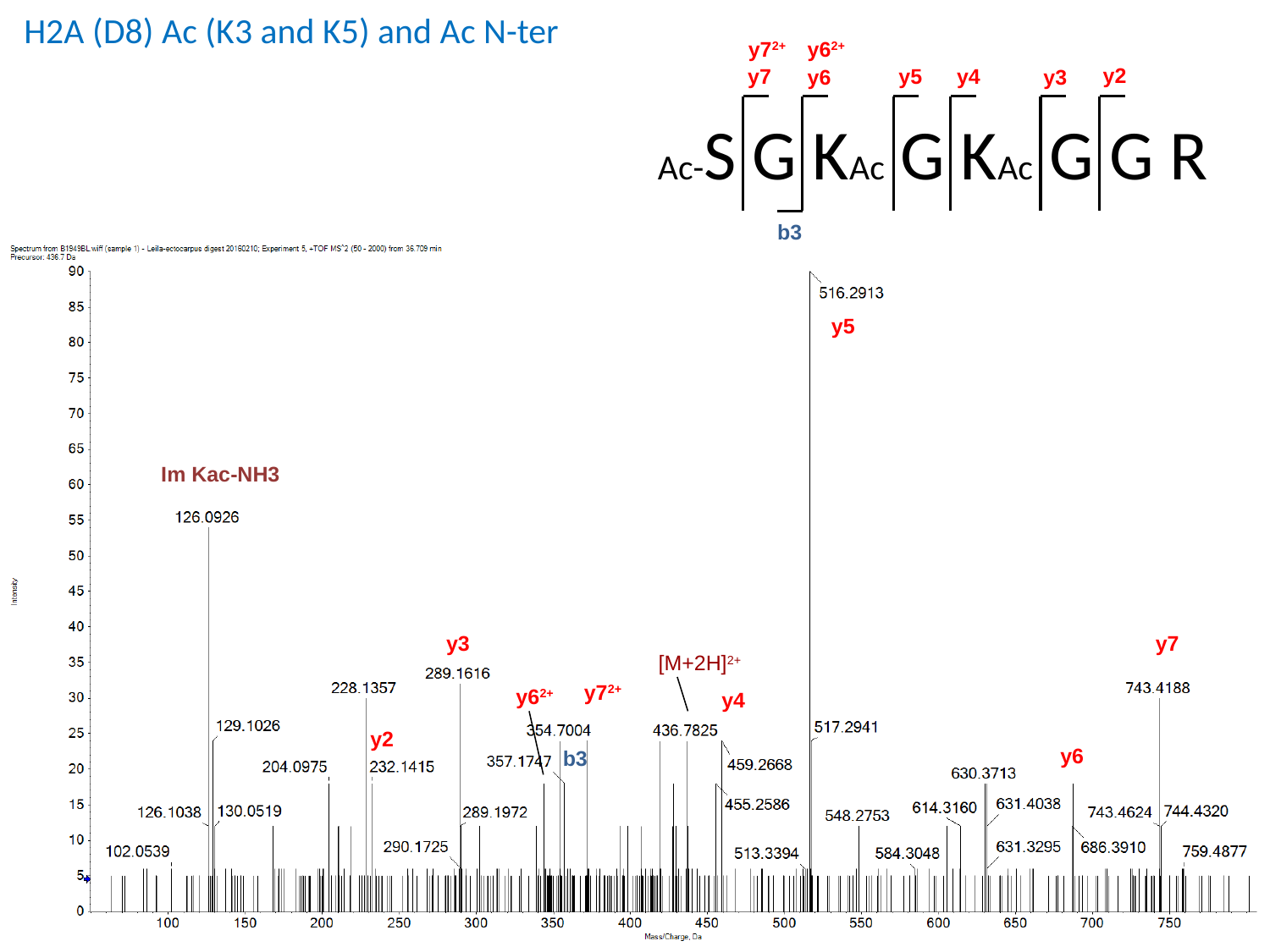

H2A (D8) Ac (K3 and K5) and Ac N-ter
y7^2+^
y6^2+^
y2
y7
y5
y4
y6
y3
Ac-S G KAc G KAc G G R
b3
y5
Im Kac-NH3
y3
y7
[M+2H]^2+^
y7^2+^
y6^2+^
y4
y2
y6
b3

### Slide 9
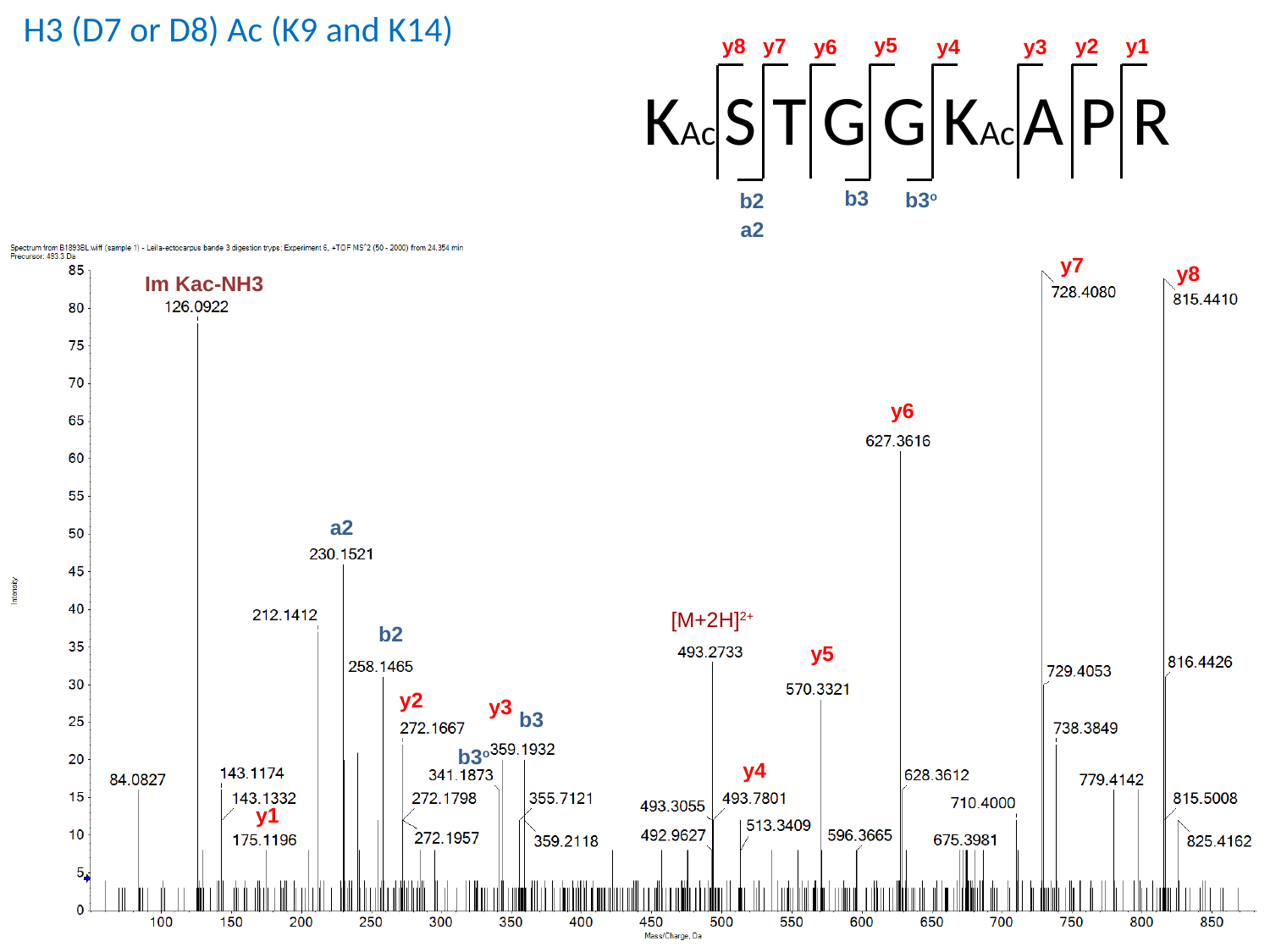

H3 (D7 or D8) Ac (K9 and K14)
y5
y7
y2
y1
y8
y4
y6
y3
KAc S T G G KAc A P R
b3
b3^o^
b2
a2
y7
y8
Im Kac-NH3
y6
a2
[M+2H]^2+^
b2
y5
y2
y3
b3
b3^o^
y4
y1

### Slide 10
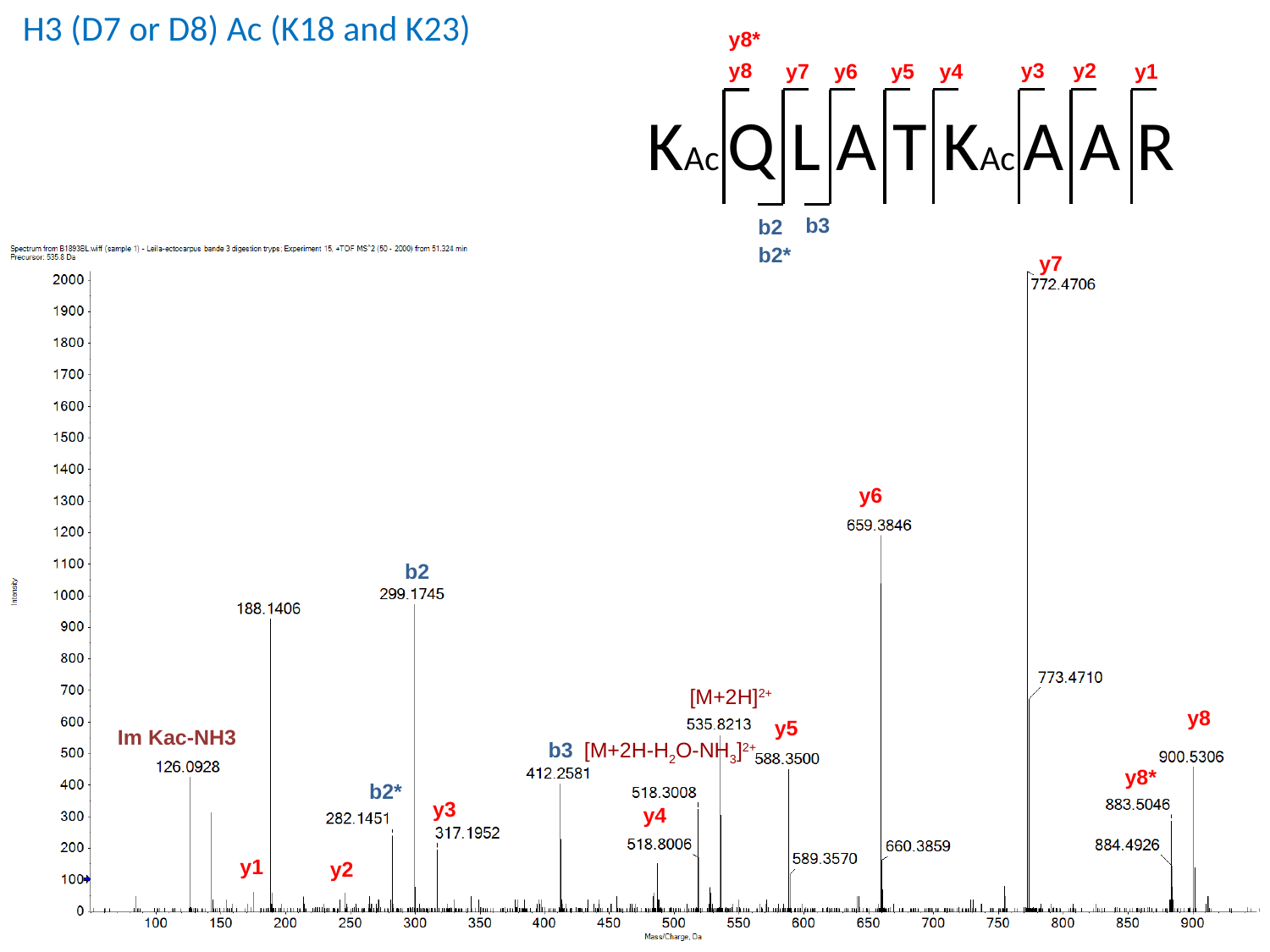

H3 (D7 or D8) Ac (K18 and K23)
y8*
y8
y3
y2
y4
y7
y5
y6
y1
KAc Q L A T KAc A A R
b3
b2
b2*
y7
y6
b2
[M+2H]^2+^
y8
y5
Im Kac-NH3
b3
[M+2H-H_2_O-NH_3_]^2+^
y8*
b2*
y3
y4
y1
y2

### Slide 11
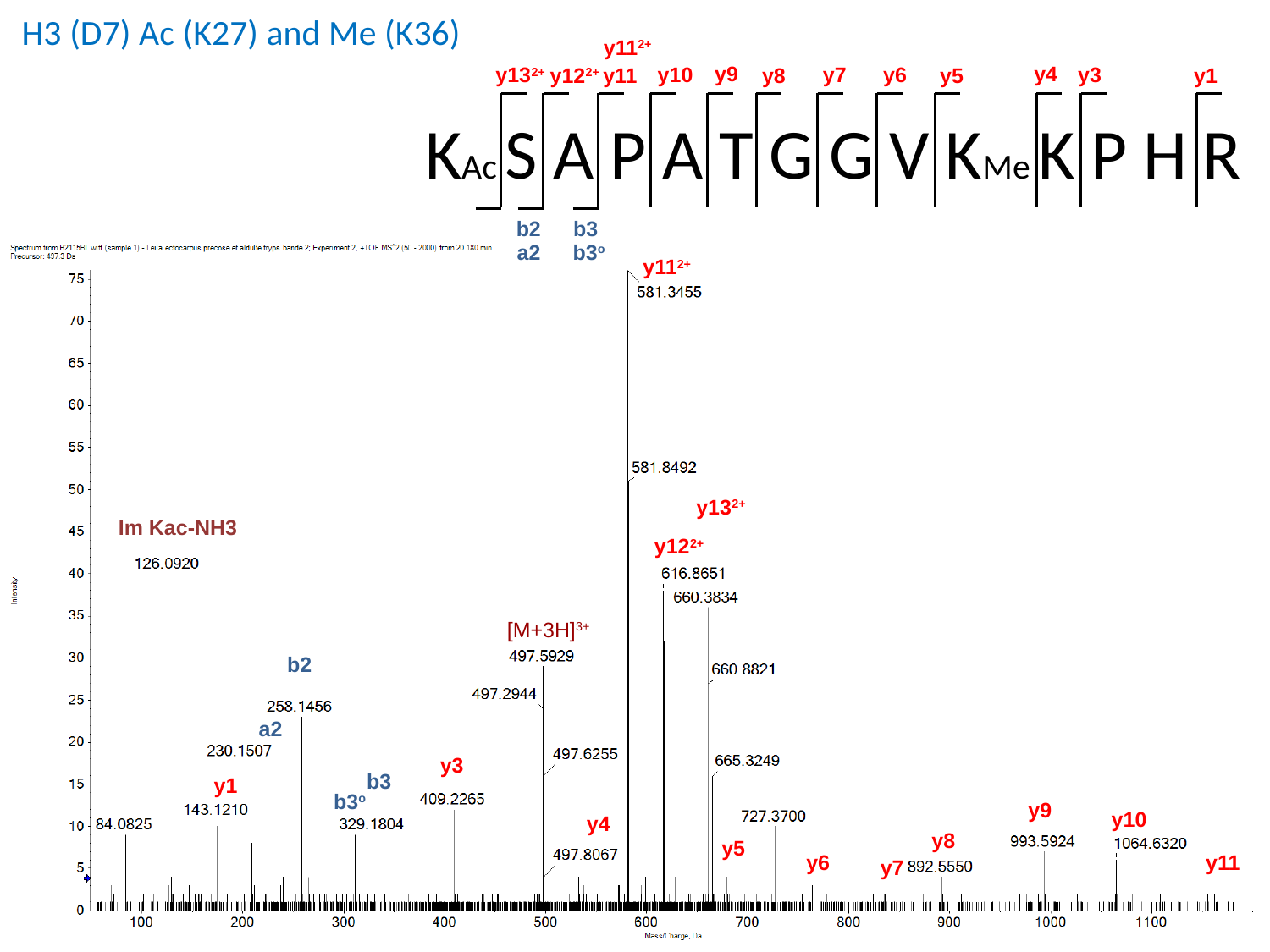

H3 (D7) Ac (K27) and Me (K36)
y11^2+^
y9
y4
y3
y6
y10
y13^2+^
y7
y12^2+^
y11
y5
y8
y1
KAc S A P A T G G V KMe K P H R
b3
b2
a2
b3^o^
y11^2+^
y13^2+^
Im Kac-NH3
y12^2+^
[M+3H]^3+^
b2
a2
y3
b3
y1
b3^o^
y9
y10
y4
y8
y5
y11
y6
y7

### Slide 12
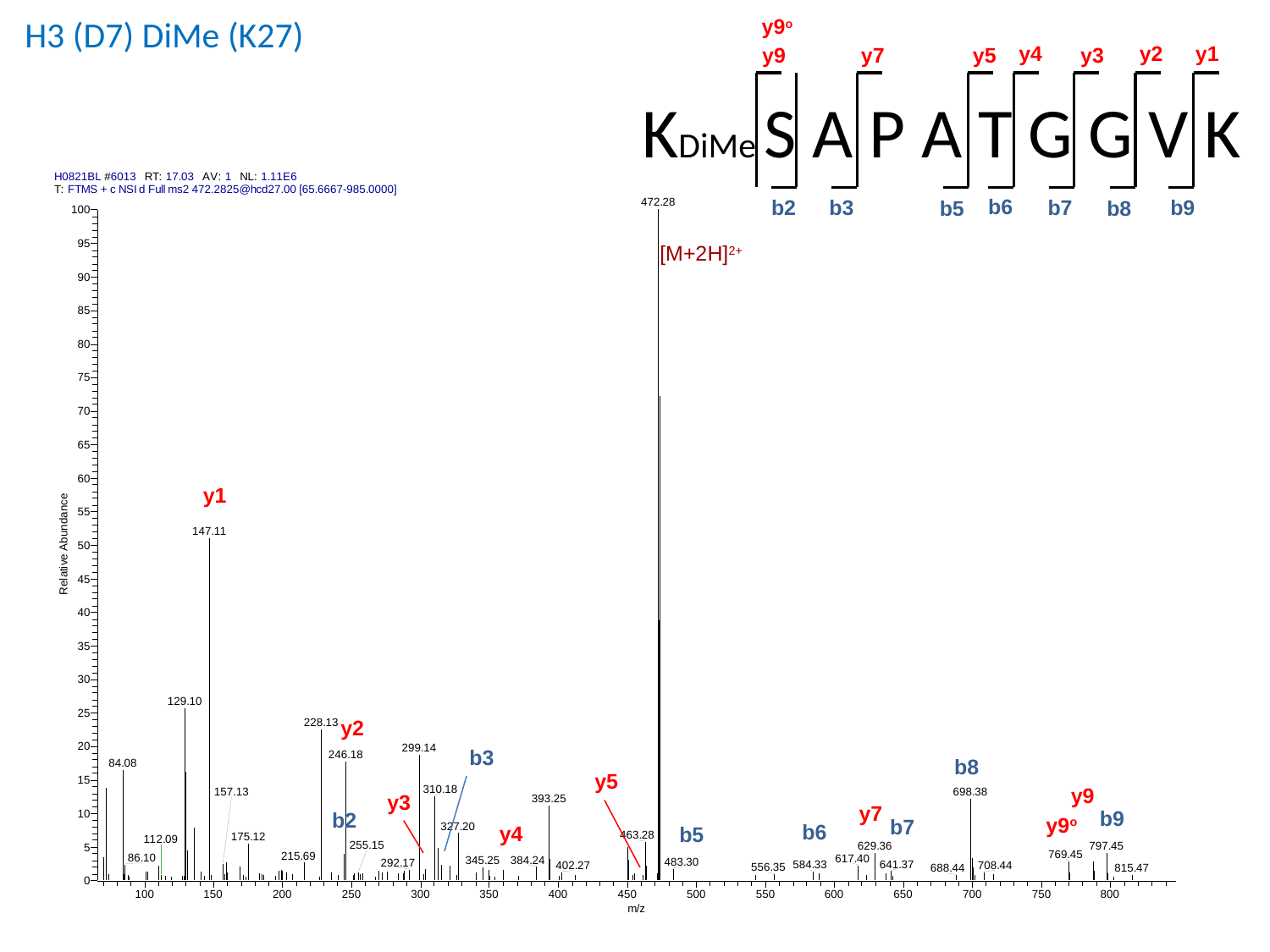

H3 (D7) DiMe (K27)
y9^o^
y4
y2
y1
y9
y5
y3
y7
KDiMe S A P A T G G V K
b6
b7
b2
b9
b3
b5
b8
[M+2H]^2+^
y1
y2
b3
b8
y5
y9
y3
y7
b9
b2
y9^o^
b7
b6
y4
b5

### Slide 13
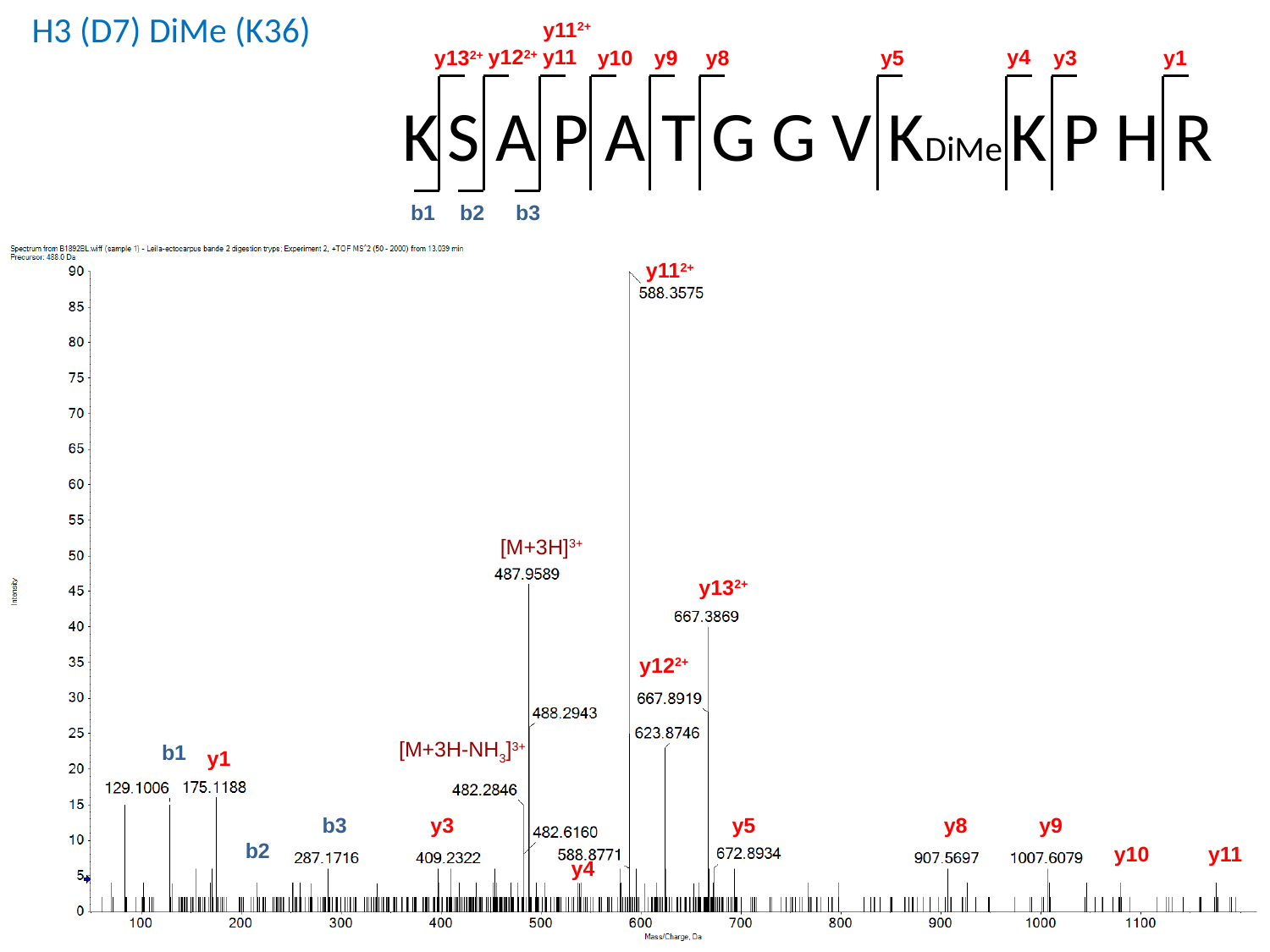

H3 (D7) DiMe (K36)
y11^2+^
y11
y4
y12^2+^
y9
y3
y1
y13^2+^
y10
y8
y5
K S A P A T G G V KDiMe K P H R
b2
b1
b3
y11^2+^
[M+3H]^3+^
y13^2+^
y12^2+^
[M+3H-NH_3_]^3+^
b1
y1
b3
y9
y8
y5
y3
b2
y10
y11
y4

### Slide 14
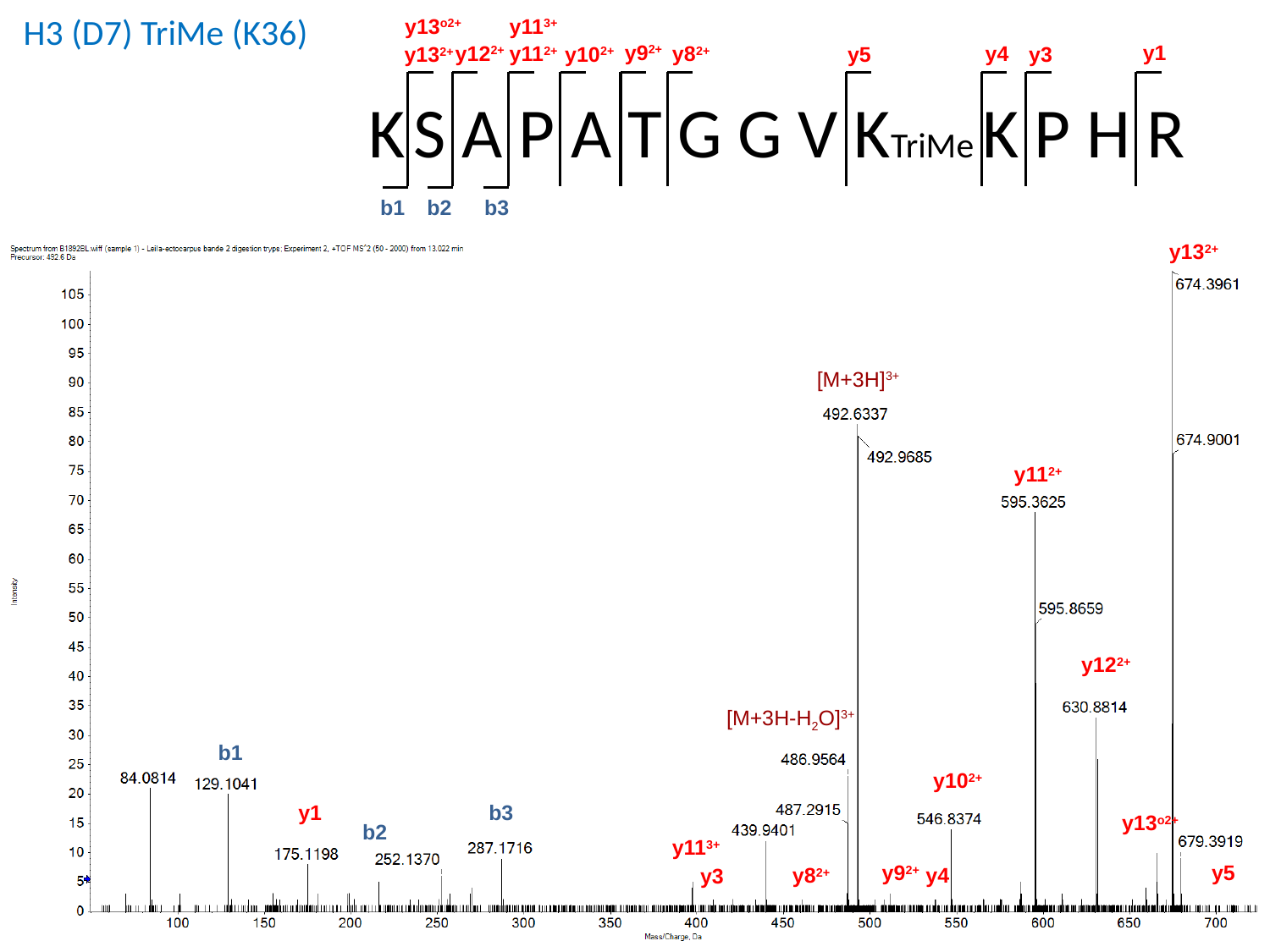

H3 (D7) TriMe (K36)
y13^o2+^
y11^3+^
y9^2+^
y1
y12^2+^
y4
y8^2+^
y11^2+^
y10^2+^
y5
y3
y13^2+^
K S A P A T G G V KTriMe K P H R
b2
b3
b1
y13^2+^
[M+3H]^3+^
y11^2+^
y12^2+^
[M+3H-H_2_O]^3+^
b1
y10^2+^
b3
y1
y13^o2+^
b2
y11^3+^
y9^2+^
y5
y4
y8^2+^
y3

### Slide 15
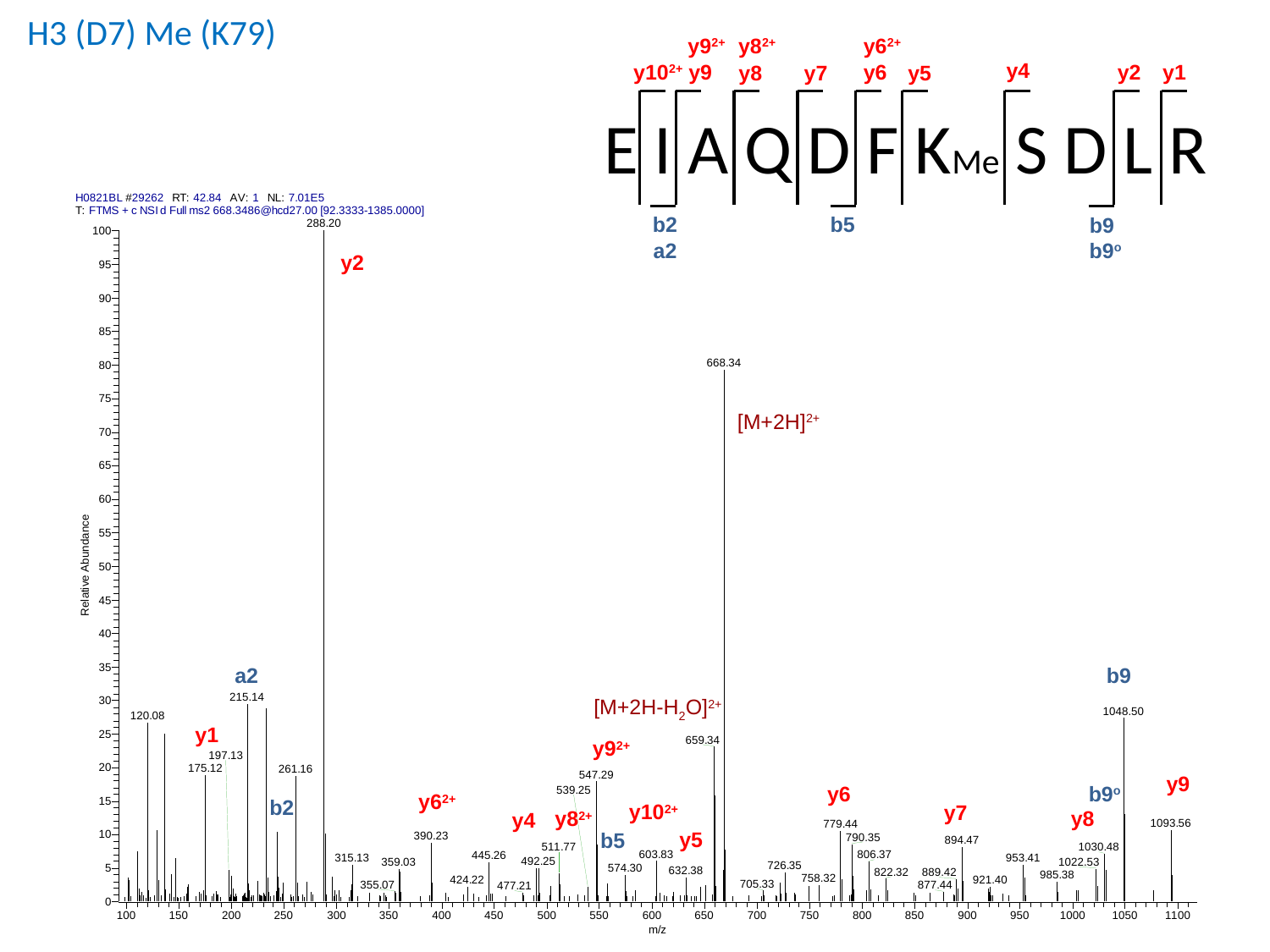

H3 (D7) Me (K79)
y9^2+^
y8^2+^
y6^2+^
y4
y10^2+^
y9
y1
y2
y6
y7
y5
y8
E I A Q D F KMe S D L R
b2
b5
b9
a2
b9^o^
y2
[M+2H]^2+^
b9
a2
[M+2H-H_2_O]^2+^
y1
y9^2+^
y9
b9^o^
y6
y6^2+^
b2
y10^2+^
y7
y8^2+^
y8
y4
y5
b5

### Slide 16
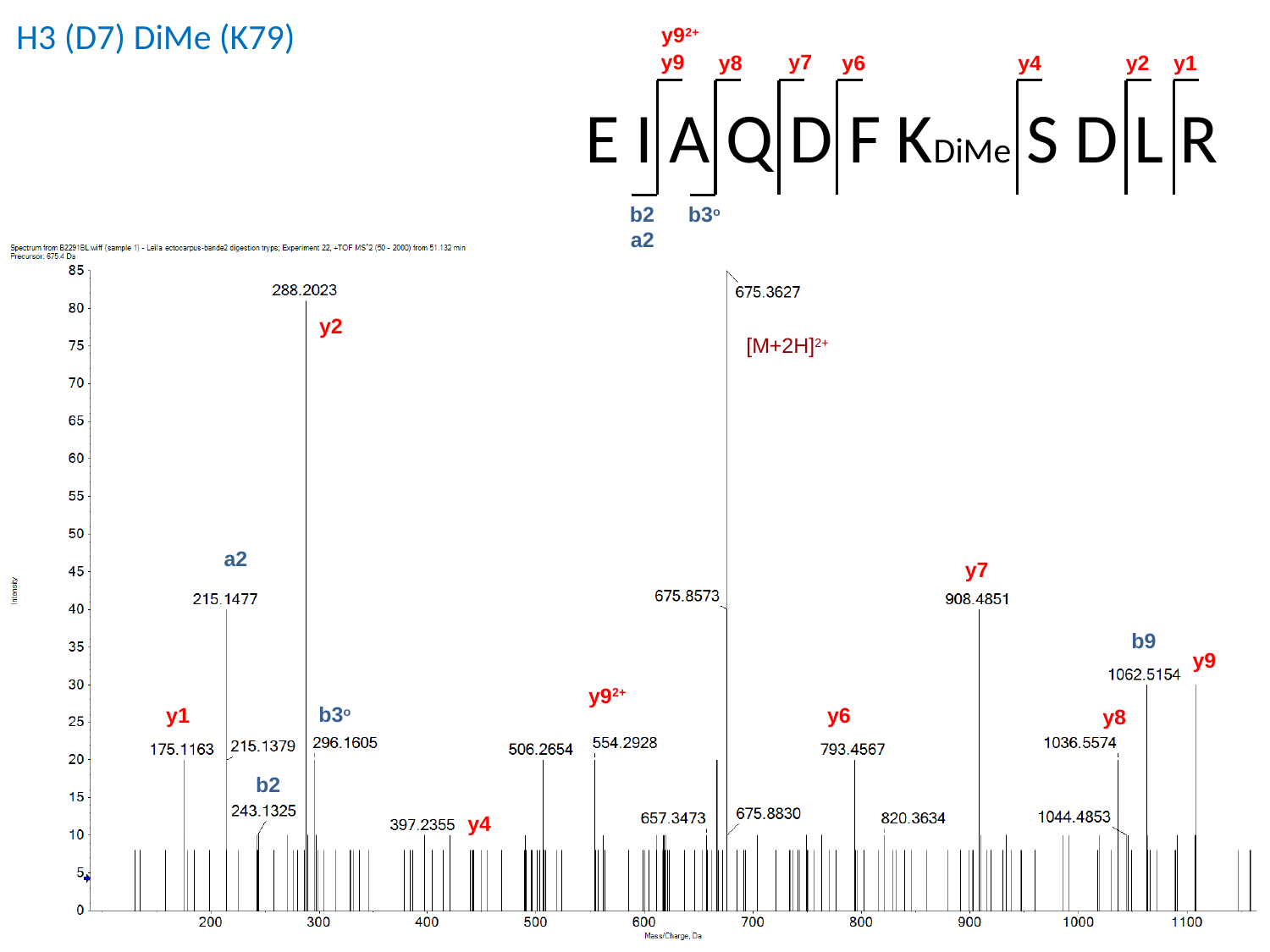

H3 (D7) DiMe (K79)
y9^2+^
y9
y7
y6
y1
y2
y4
y8
E I A Q D F KDiMe S D L R
b2
b3^o^
a2
y2
[M+2H]^2+^
a2
y7
b9
y9
y9^2+^
b3^o^
y6
y1
y8
b2
y4

### Slide 17
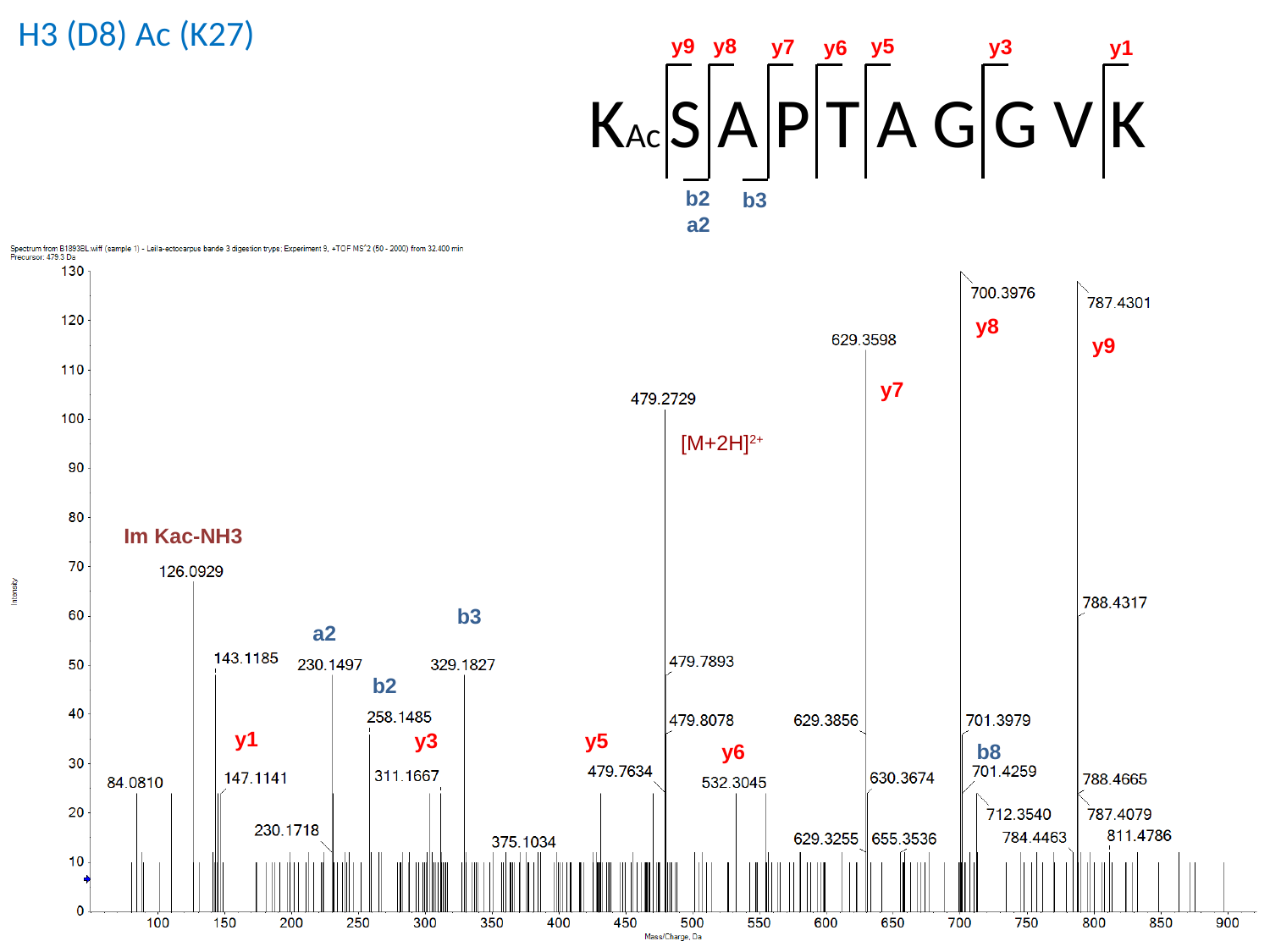

H3 (D8) Ac (K27)
y8
y5
y9
y7
y3
y6
y1
KAc S A P T A G G V K
b2
b3
a2
y8
y9
y7
[M+2H]^2+^
Im Kac-NH3
b3
a2
b2
y1
y5
y3
b8
y6

### Slide 18
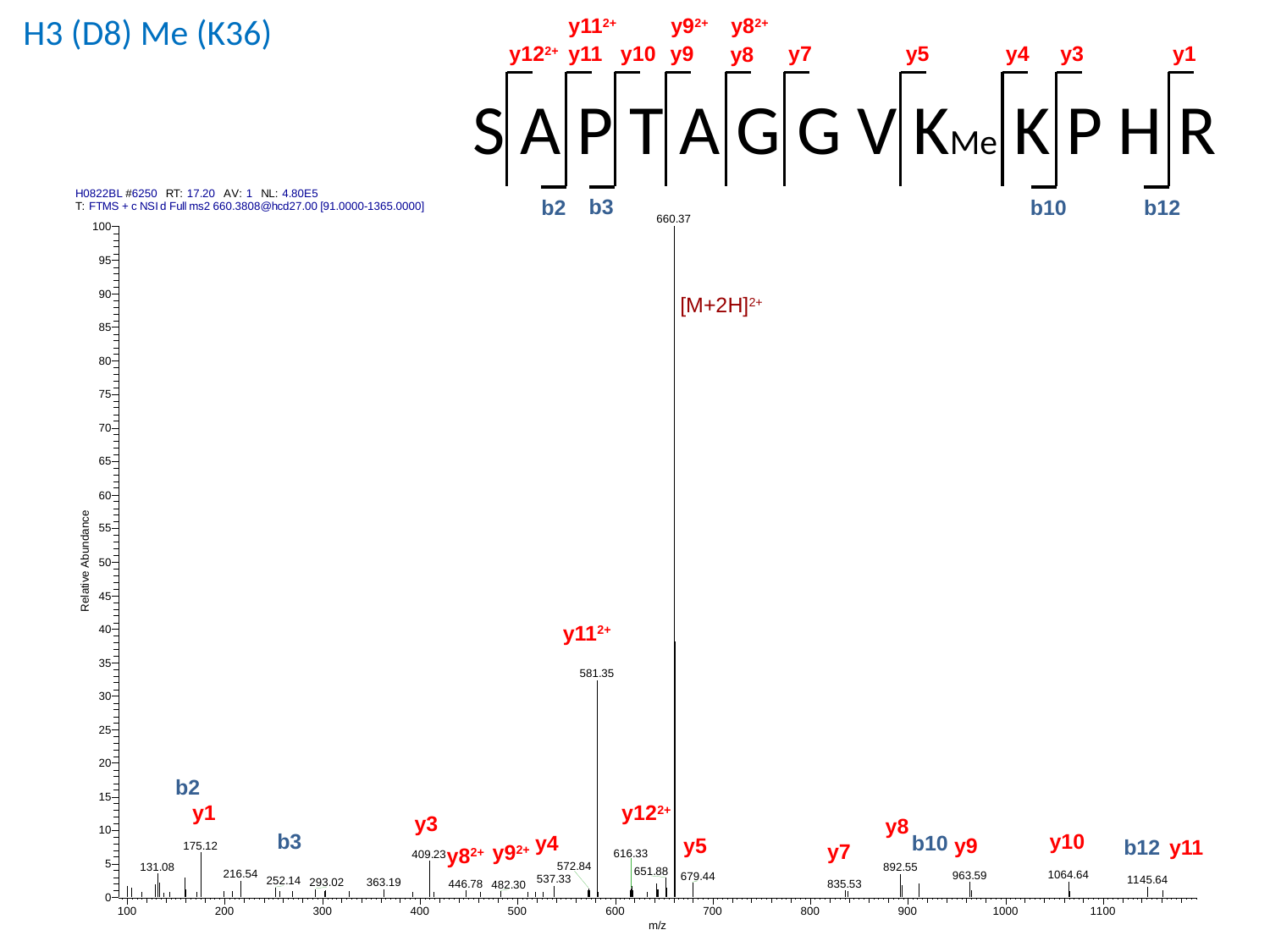

H3 (D8) Me (K36)
y11^2+^
y9^2+^
y8^2+^
y11
y10
y7
y3
y1
y5
y4
y12^2+^
y9
y8
S A P T A G G V KMe K P H R
b3
b10
b2
b12
[M+2H]^2+^
y11^2+^
b2
y1
y12^2+^
y3
y8
b3
y10
y4
b10
y5
y9
b12
y11
y7
y9^2+^
y8^2+^

### Slide 19
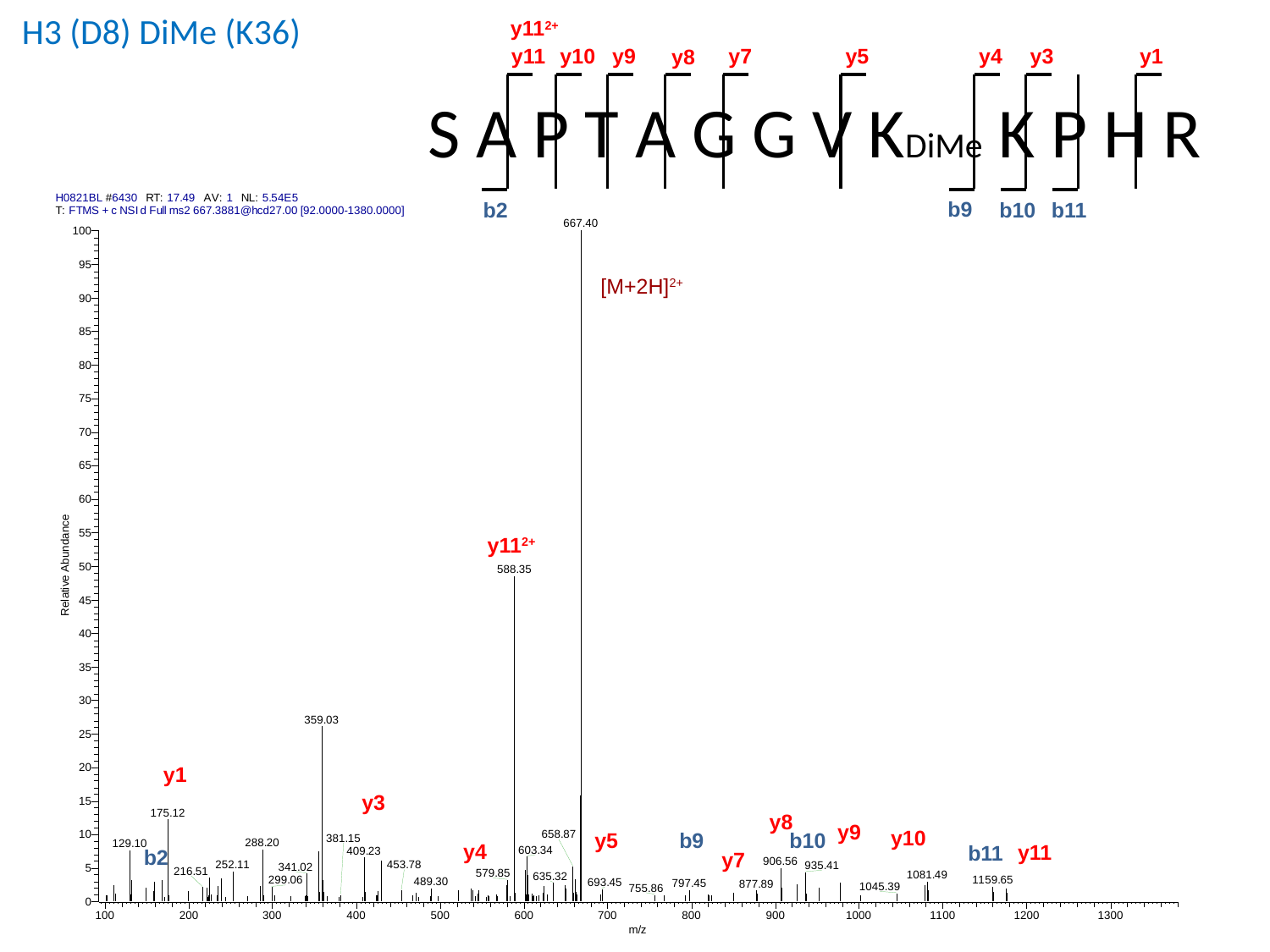

H3 (D8) DiMe (K36)
y11^2+^
y11
y4
y3
y10
y7
y5
y9
y1
y8
S A P T A G G V KDiMe K P H R
b9
b2
b10
b11
[M+2H]^2+^
y11^2+^
y1
y3
y8
y9
y10
b9
y5
b10
y4
y11
b11
b2
y7

### Slide 20
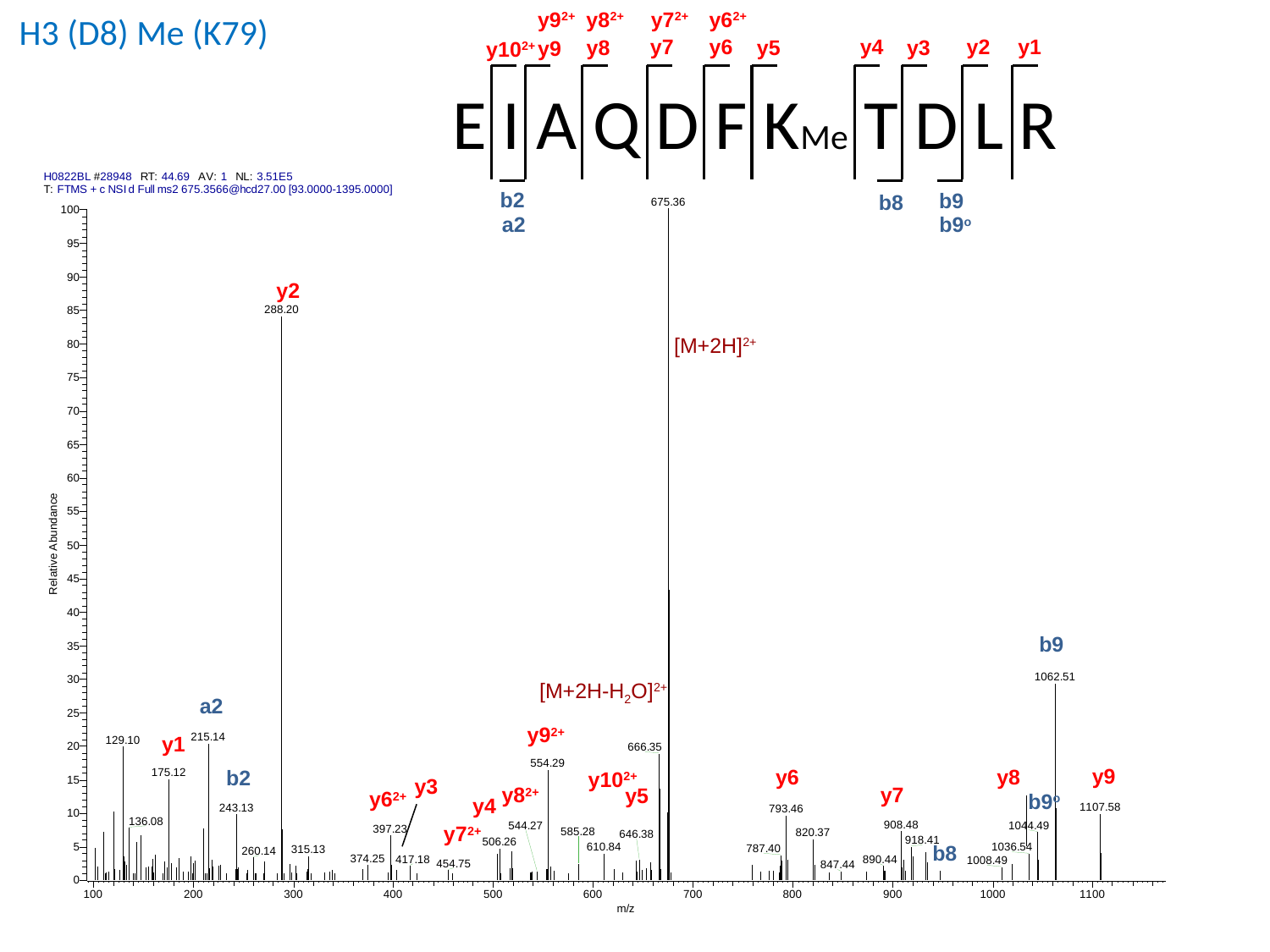

y9^2+^
y8^2+^
y7^2+^
y6^2+^
y7
y6
y4
y1
y2
y8
y5
y3
y9
y10^2+^
E I A Q D F KMe T D L R
b2
b9
b8
a2
b9^o^
H3 (D8) Me (K79)
y2
[M+2H]^2+^
b9
[M+2H-H_2_O]^2+^
a2
y9^2+^
y1
y9
y8
y6
b2
y10^2+^
y3
y8^2+^
y7
y5
y6^2+^
b9^o^
y4
y7^2+^
b8

### Slide 21
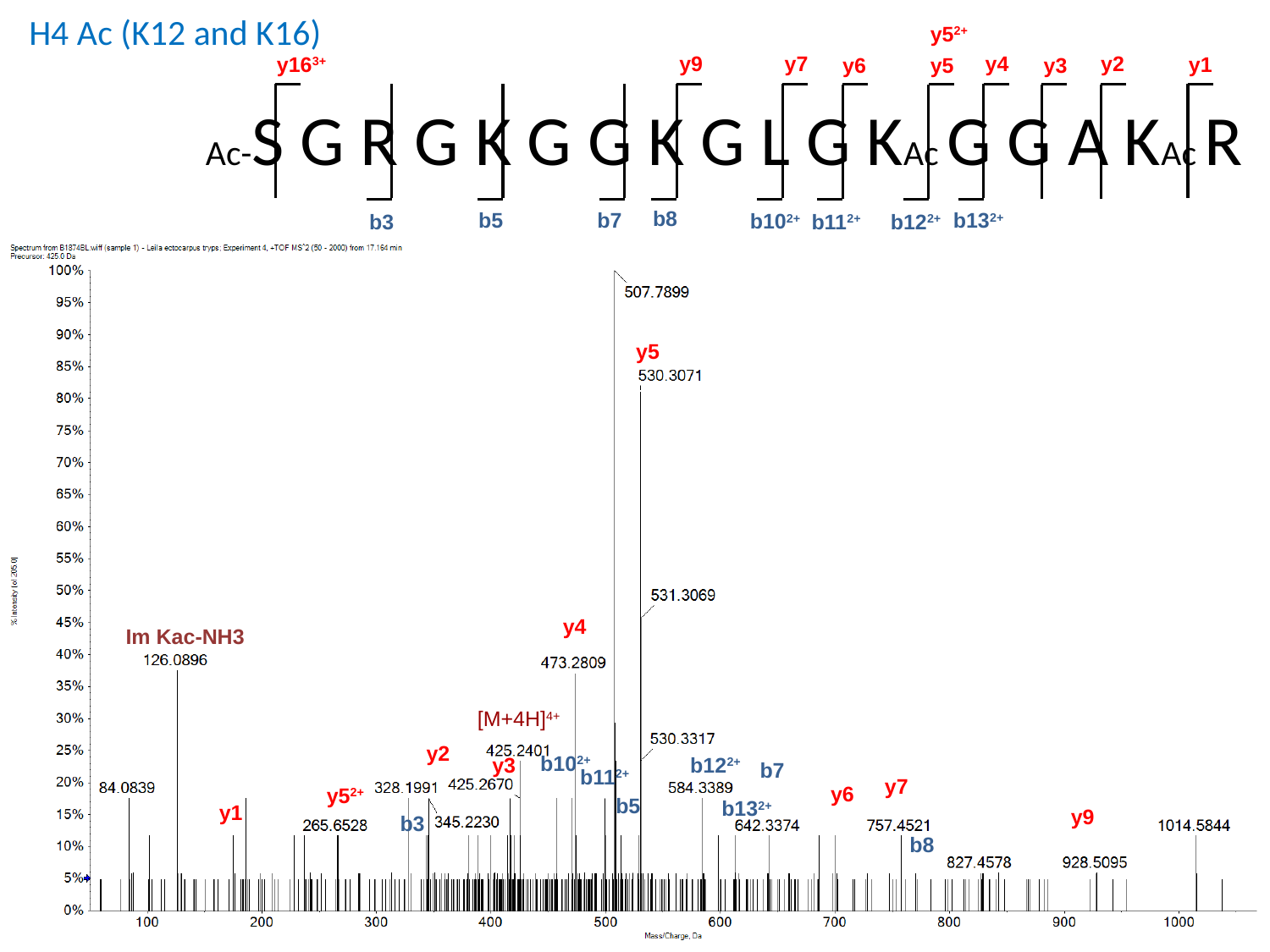

H4 Ac (K12 and K16)
y5^2+^
y4
y2
y7
y9
y16^3+^
y1
y6
y5
y3
Ac-S G R G K G G K G L G KAc G G A KAc R
b8
b7
b5
b13^2+^
b10^2+^
b3
b11^2+^
b12^2+^
y5
y4
[M+4H]^4+^
y2
b10^2+^
y3
b12^2+^
b7
b11^2+^
y7
y6
y5^2+^
b5
b13^2+^
y1
y9
b3
b8
Im Kac-NH3

### Slide 22
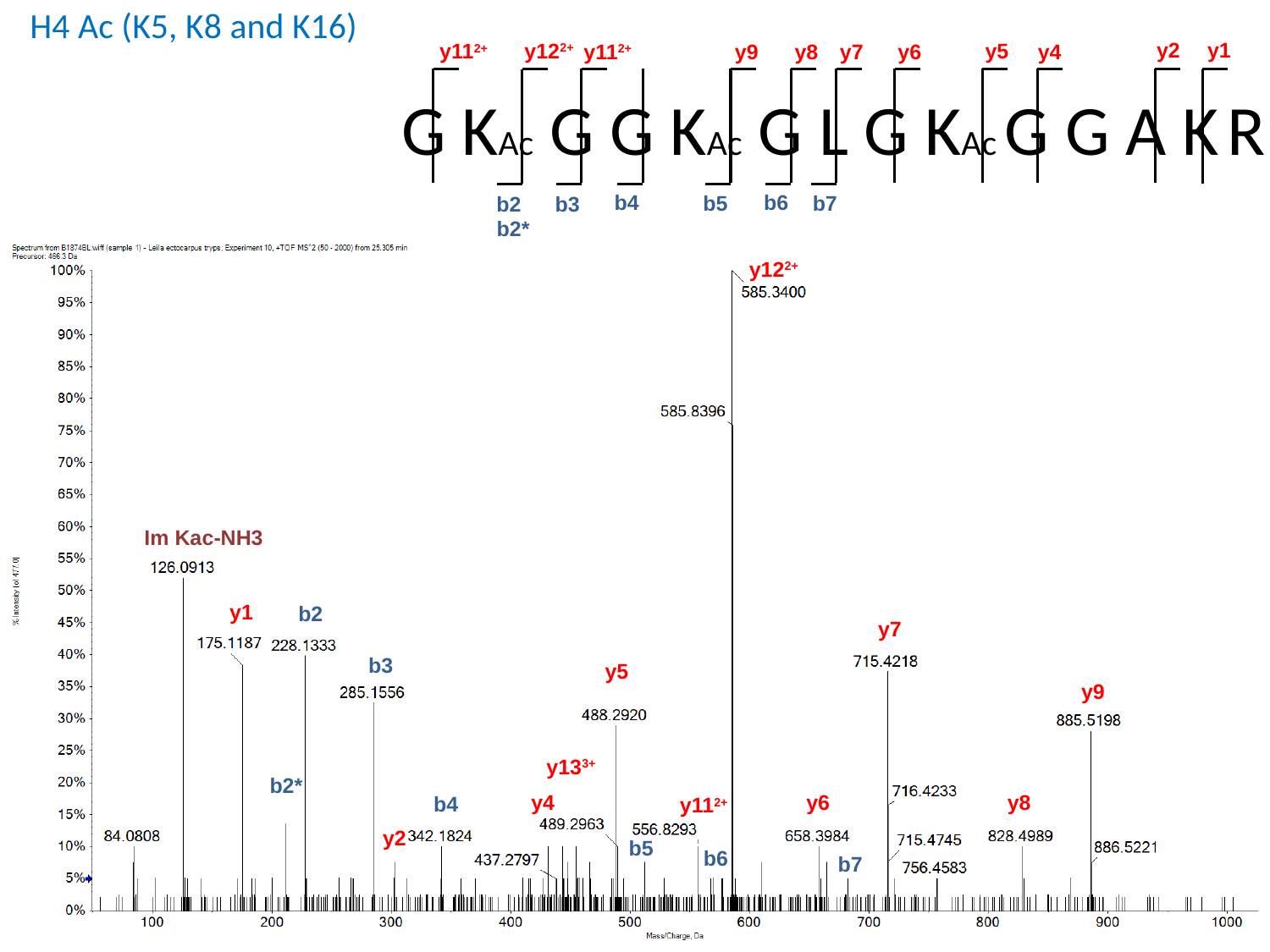

H4 Ac (K5, K8 and K16)
y2
y1
y12^2+^
y11^2+^
y5
y11^2+^
y4
y9
y8
y7
y6
G KAc G G KAc G L G KAc G G A K R
b6
b4
b7
b5
b2
b3
b2*
y12^2+^
Im Kac-NH3
y1
b2
y7
b3
y5
y9
y13^3+^
b2*
y6
y4
y8
b4
y11^2+^
y2
b5
b6
b7

### Slide 23
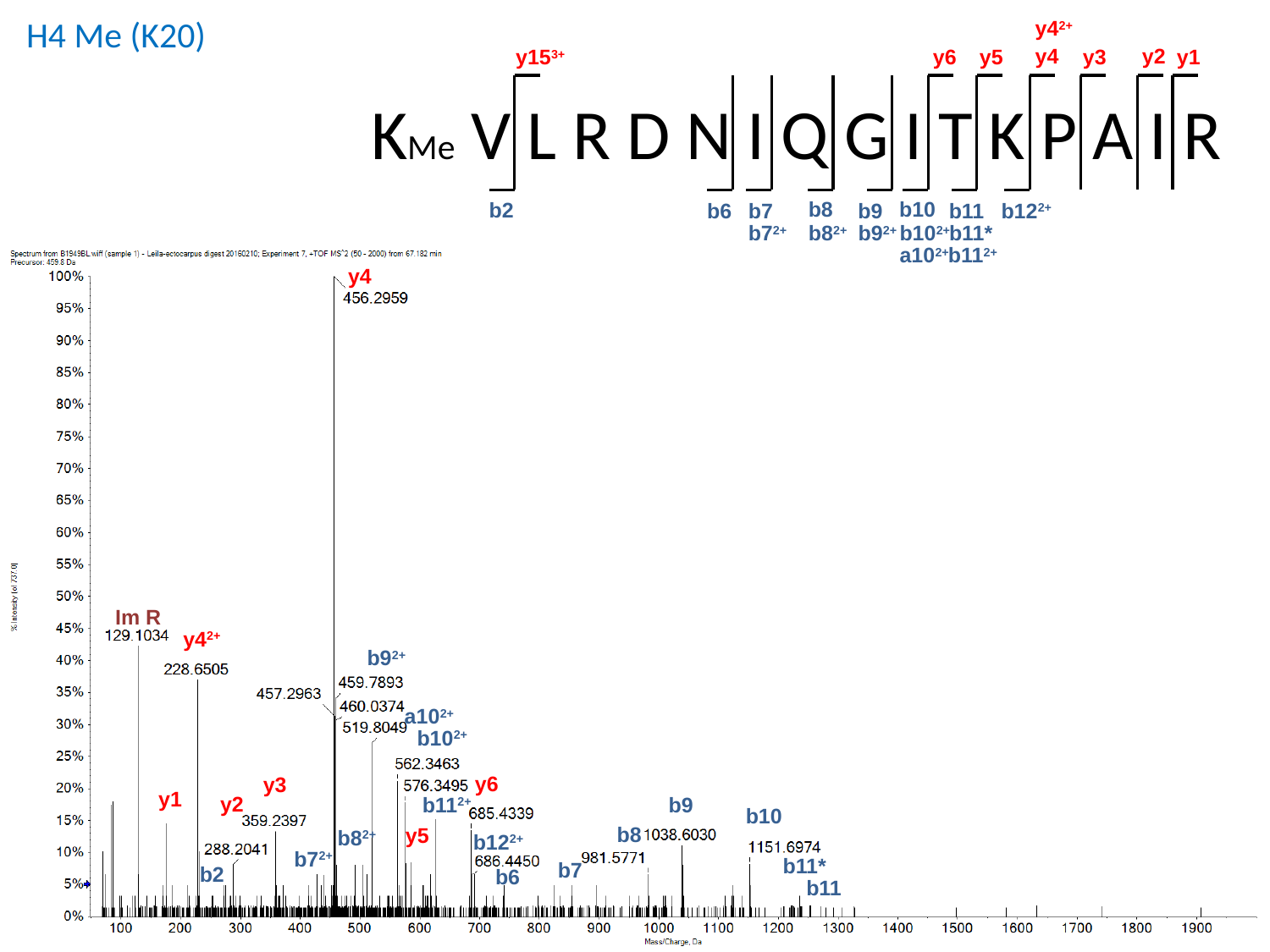

H4 Me (K20)
y4^2+^
y4
y2
y1
y5
y15^3+^
y6
y3
KMe V L R D N I Q G I T K P A I R
b10
b8
b2
b12^2+^
b11
b9
b7
b6
b7^2+^
b8^2+^
b9^2+^
b10^2+^
b11*
a10^2+^
b11^2+^
y4
y4^2+^
b9^2+^
a10^2+^
b10^2+^
y6
y3
y1
y2
b11^2+^
b9
b10
b8
y5
b8^2+^
b12^2+^
b7^2+^
b11*
b7
b2
b6
b11
Im R

### Slide 24
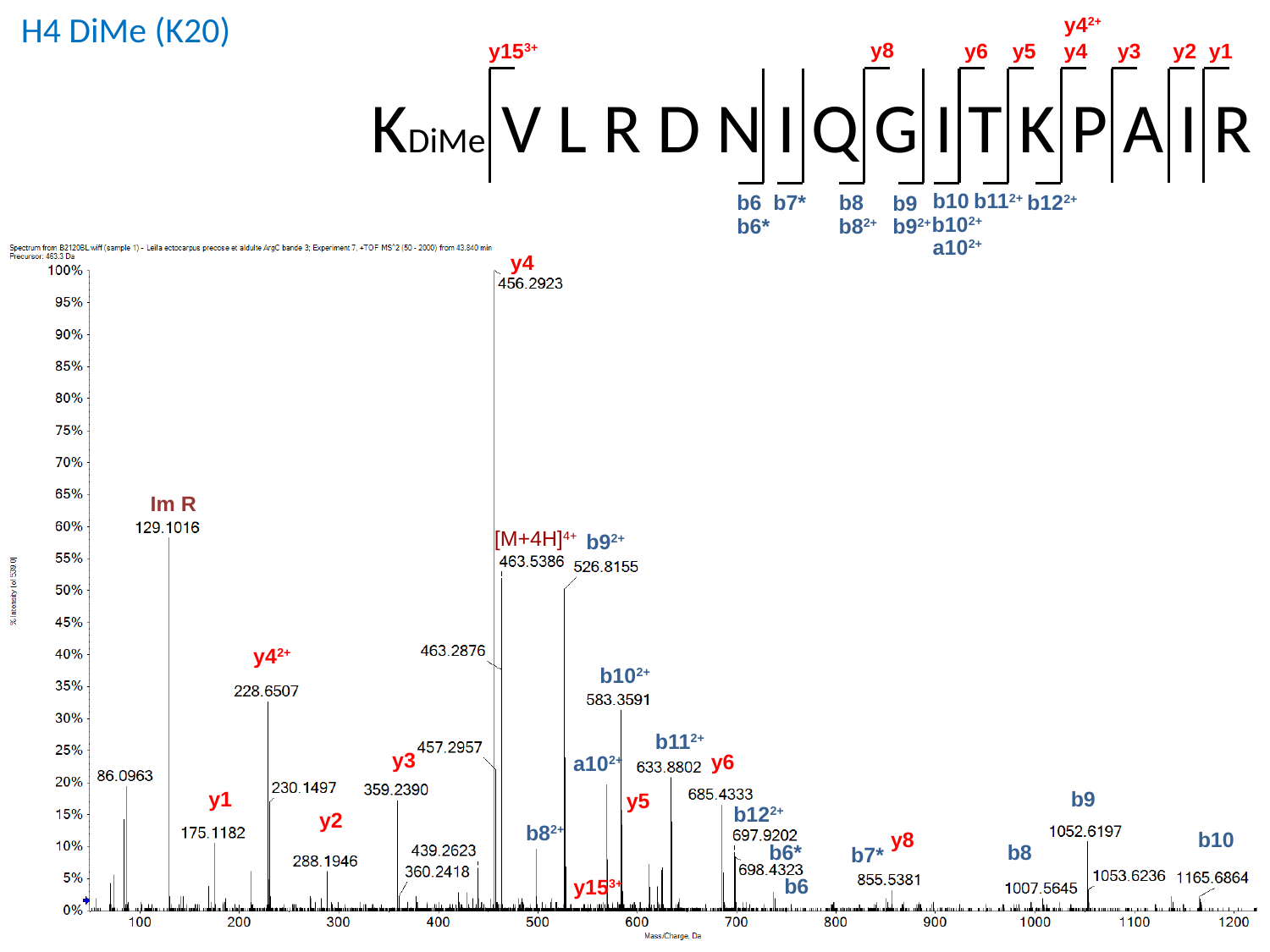

H4 DiMe (K20)
y4^2+^
y8
y15^3+^
y2
y1
y6
y3
y5
y4
KDiMe V L R D N I Q G I T K P A I R
b10
b11^2+^
b7*
b8
b12^2+^
b6
b9
b10^2+^
b6*
b8^2+^
b9^2+^
a10^2+^
[M+4H]^4+^
b9^2+^
y4^2+^
b10^2+^
b11^2+^
y3
y6
a10^2+^
b9
y1
y5
b12^2+^
y2
b8^2+^
y8
b10
b8
b6*
b7*
b6
y15^3+^
y4
Im R

### Slide 25
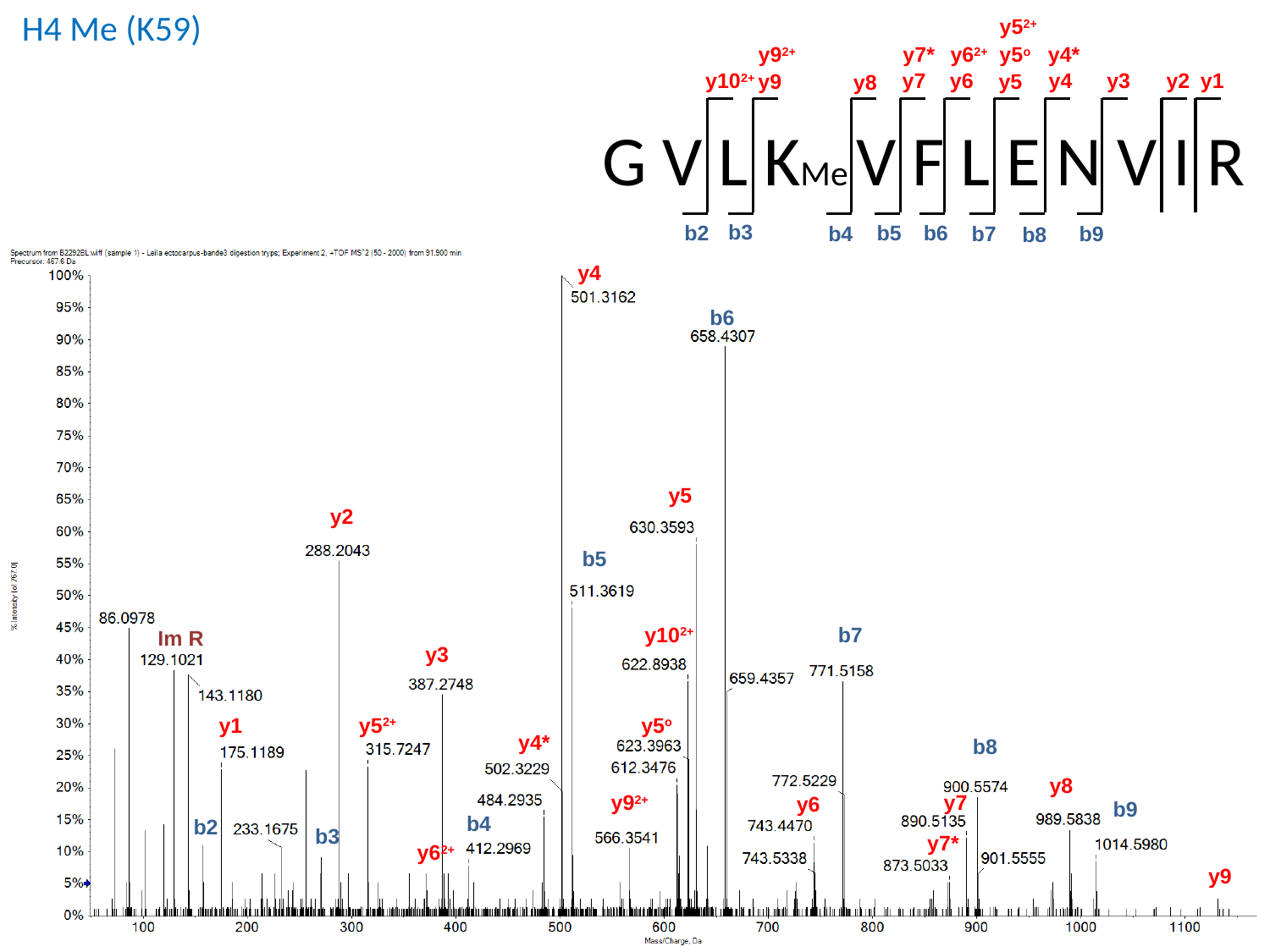

H4 Me (K59)
y5^2+^
y9^2+^
y7*
y6^2+^
y5^o^
y4*
y1
y10^2+^
y6
y3
y4
y2
y7
y9
y5
y8
G V L KMe V F L E N V I R
b3
b2
b6
b5
b4
b9
b7
b8
y4
b6
y5
y2
b5
y10^2+^
b7
y3
y5^o^
y1
y5^2+^
y4*
b8
y8
y7
y9^2+^
y6
b9
b4
b2
b3
y7*
y6^2+^
y9
Im R

### Slide 26
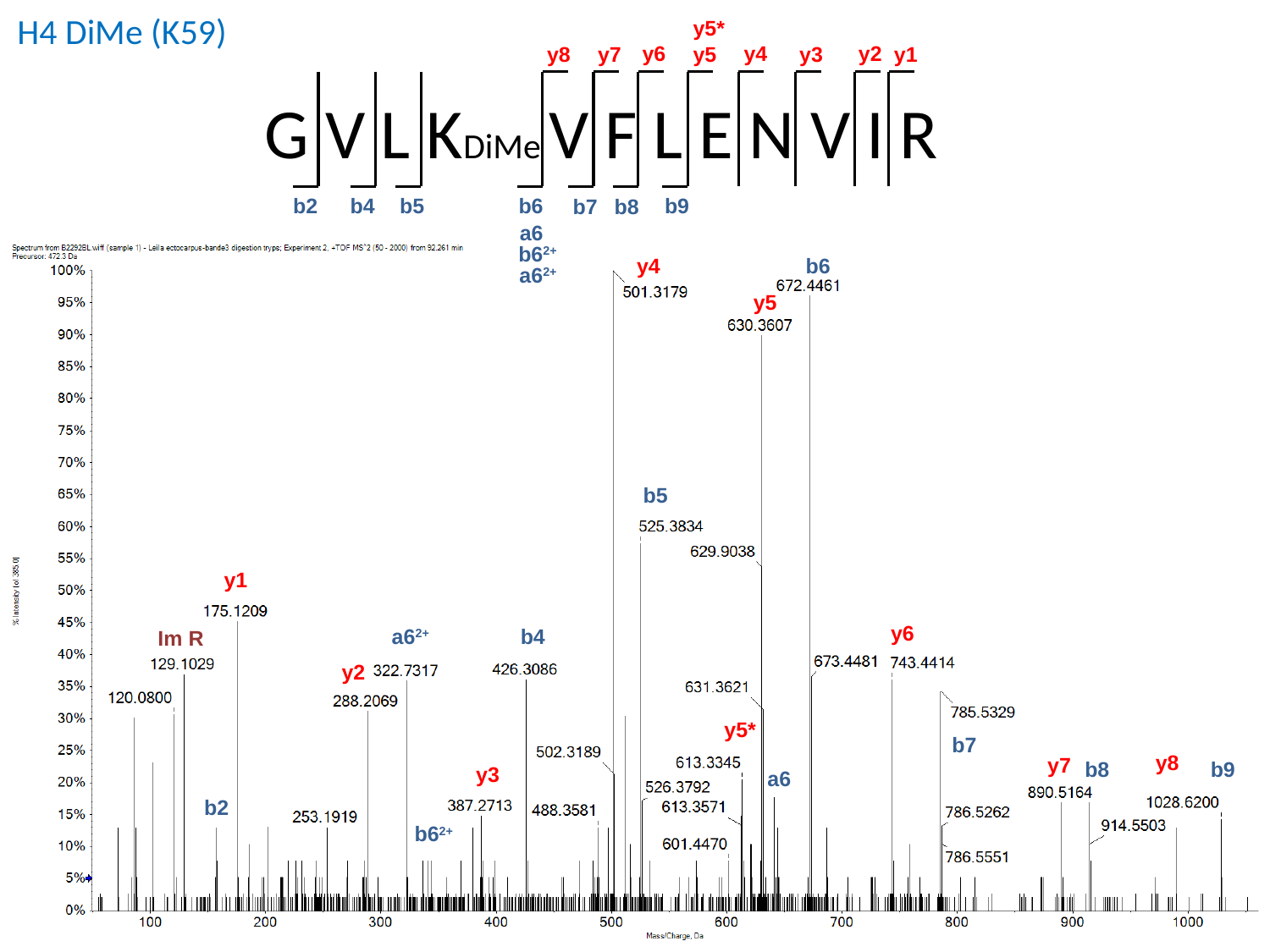

H4 DiMe (K59)
y5*
y4
y2
y6
y8
y5
y1
y7
y3
G V L KDiMe V F L E N V I R
b2
b9
b4
b5
b6
b7
b8
a6
b6^2+^
a6^2+^
b6
y5
b5
y1
y6
a6^2+^
b4
y2
y5*
b7
y8
y7
b9
b8
y3
a6
b2
b6^2+^
y4
Im R

### Slide 27
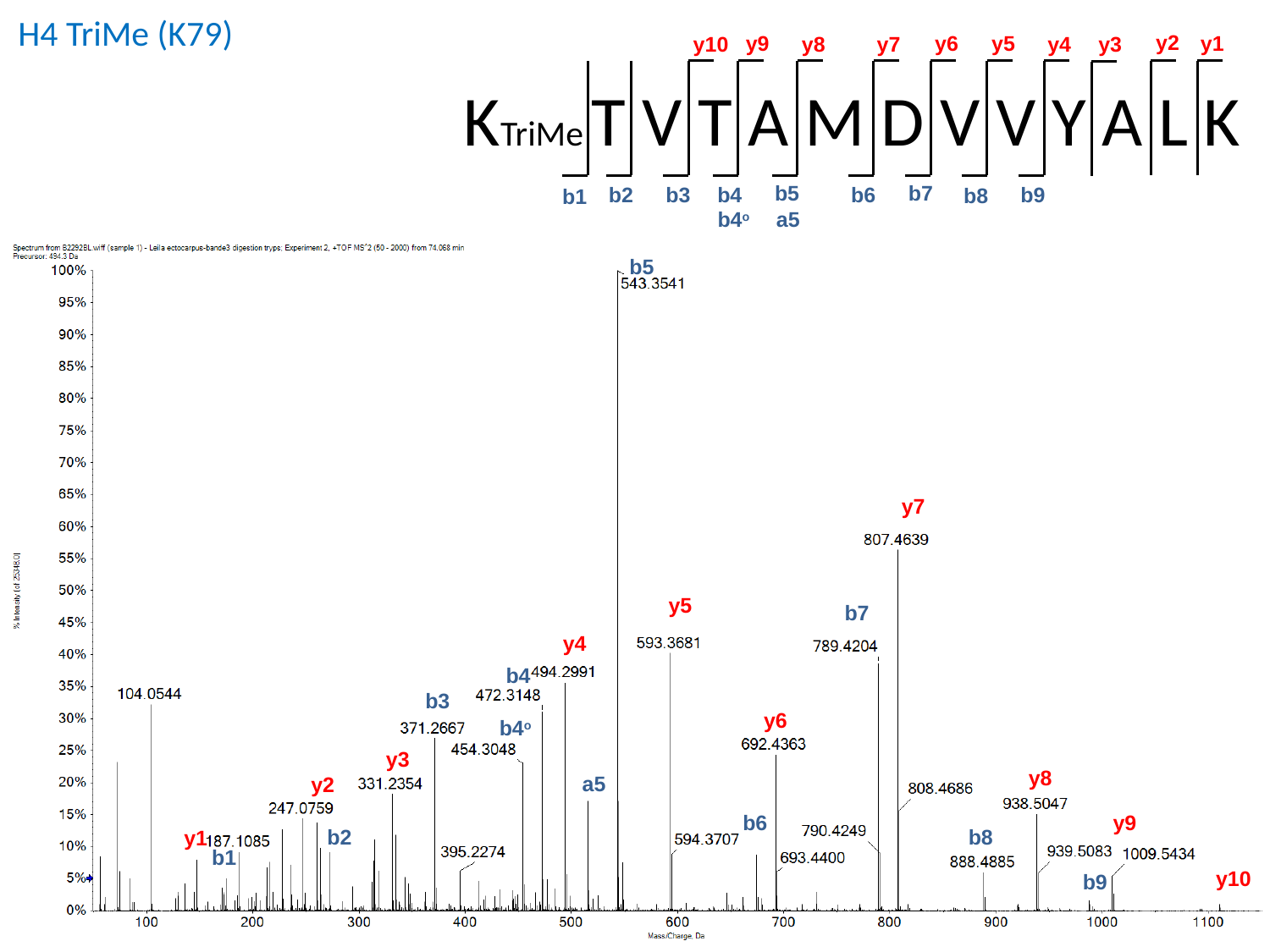

H4 TriMe (K79)
y2
y6
y5
y9
y1
y10
y4
y8
y7
y3
KTriMe T V T A M D V V Y A L K
b5
b7
b2
b6
b9
b3
b4
b8
b1
b4^o^
a5
b5
y7
y5
b7
y4
b4
b3
y6
b4^o^
y3
y8
a5
y2
b6
y9
b8
b2
y1
b1
y10
b9

### Slide 28
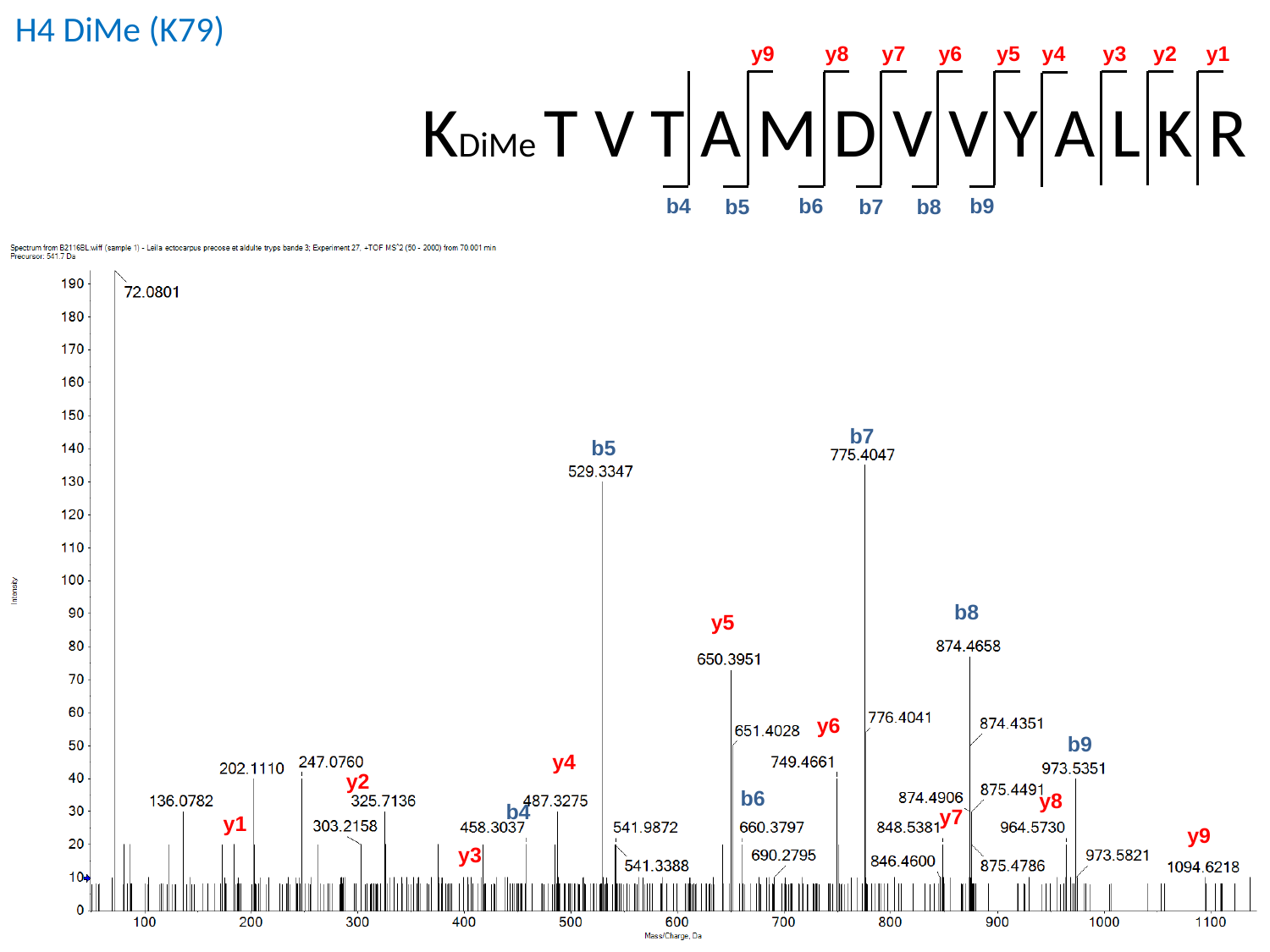

H4 DiMe (K79)
y9
y8
y7
y6
y5
y4
y3
y2
y1
KDiMe T V T A M D V V Y A L K R
b9
b6
b4
b7
b5
b8
b5
b8
y5
y6
b9
y4
y2
b6
y8
b4
y7
y1
y9
y3
b7

### Slide 29
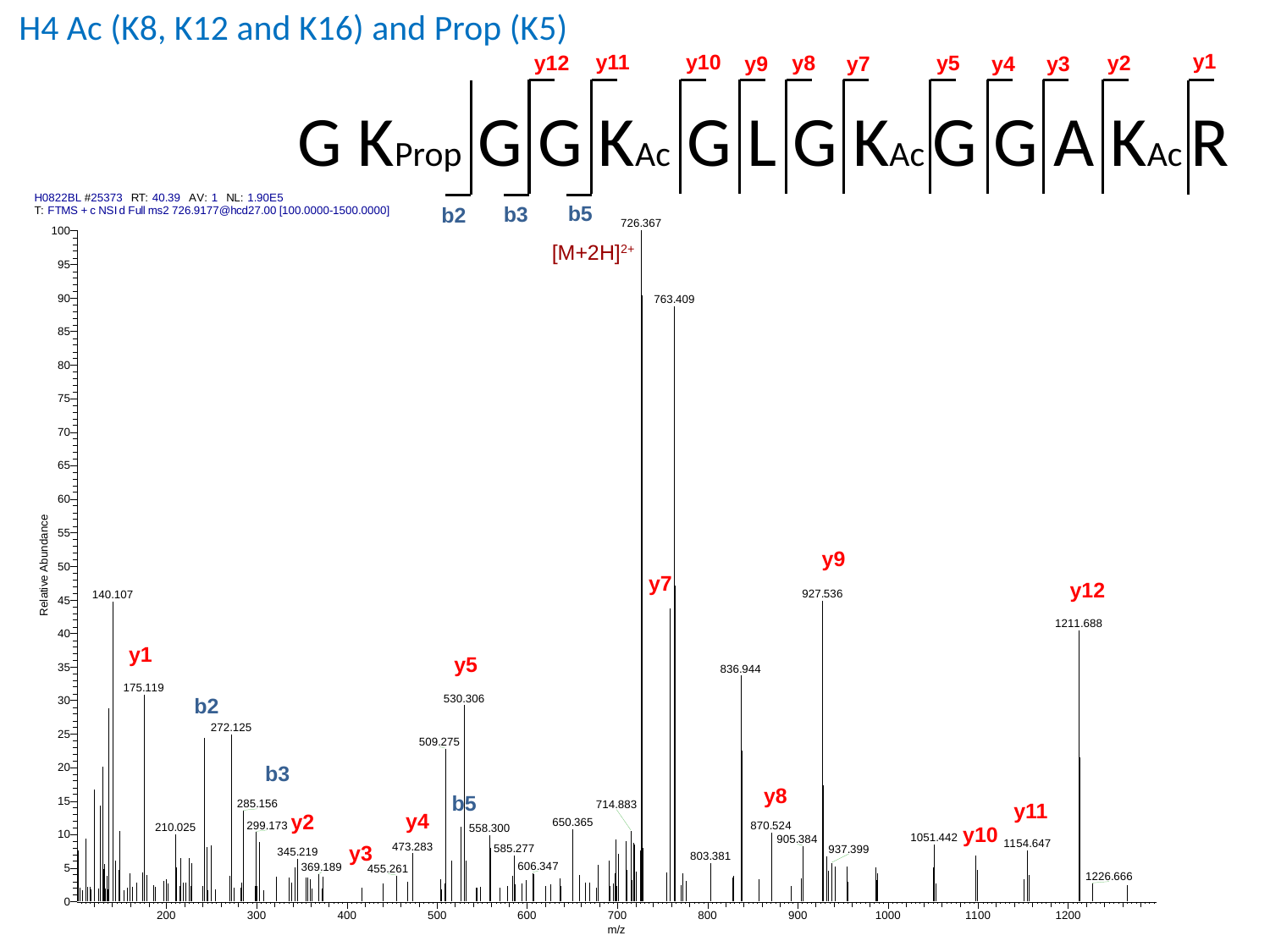

H4 Ac (K8, K12 and K16) and Prop (K5)
y1
y11
y10
y2
y12
y8
y5
y3
y9
y7
y4
G KProp G G KAc G L G KAc G G A KAc R
b5
b3
b2
[M+2H]^2+^
y9
y7
y12
y1
y5
b2
b3
y8
b5
y11
y4
y2
y10
y3
