## Supplementary material for "Histone modifications during the life cycle of the brown alga *Ectocarpus*": Figs S3 to S10

### Slide 1
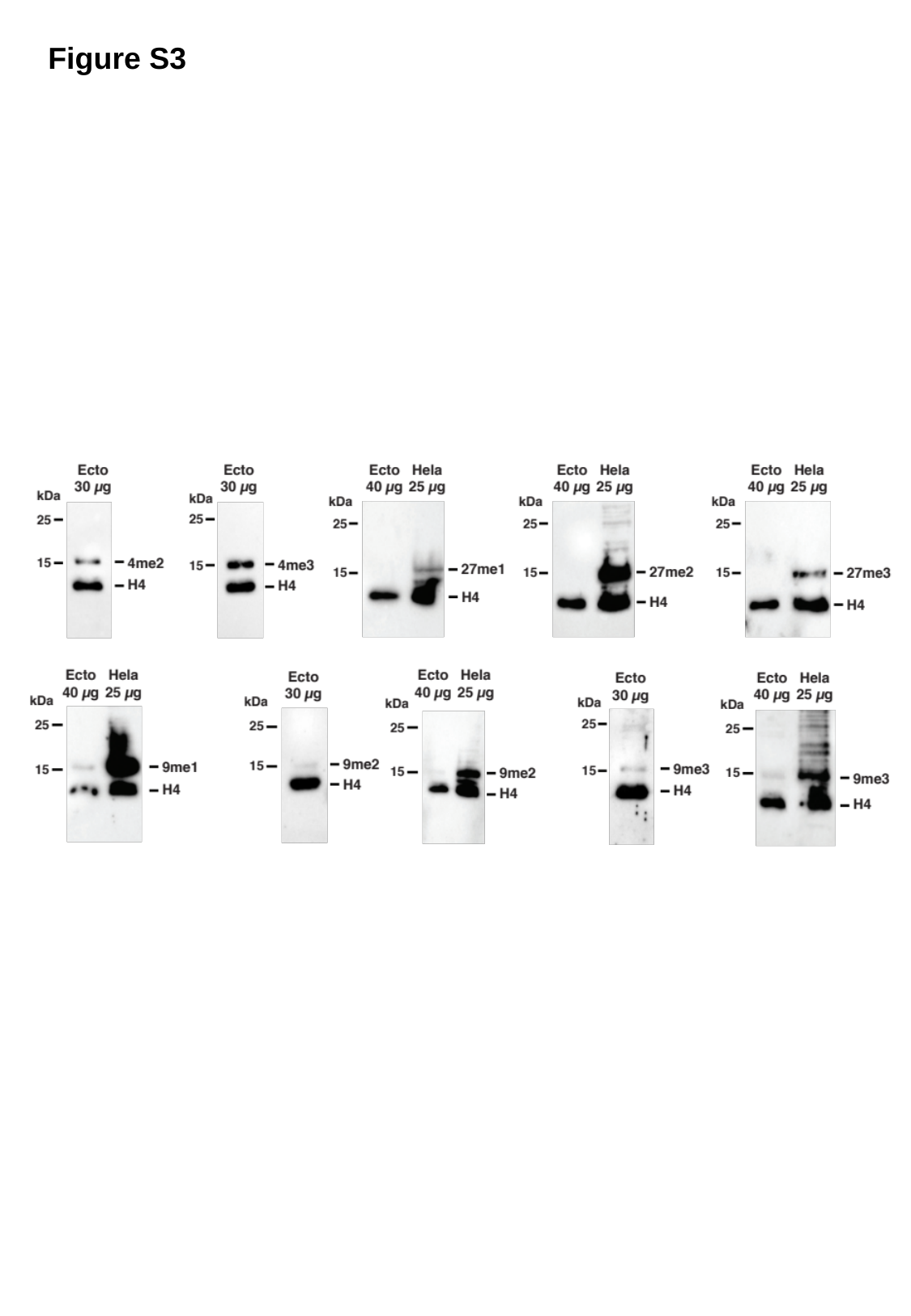

Figure S3

### Slide 2
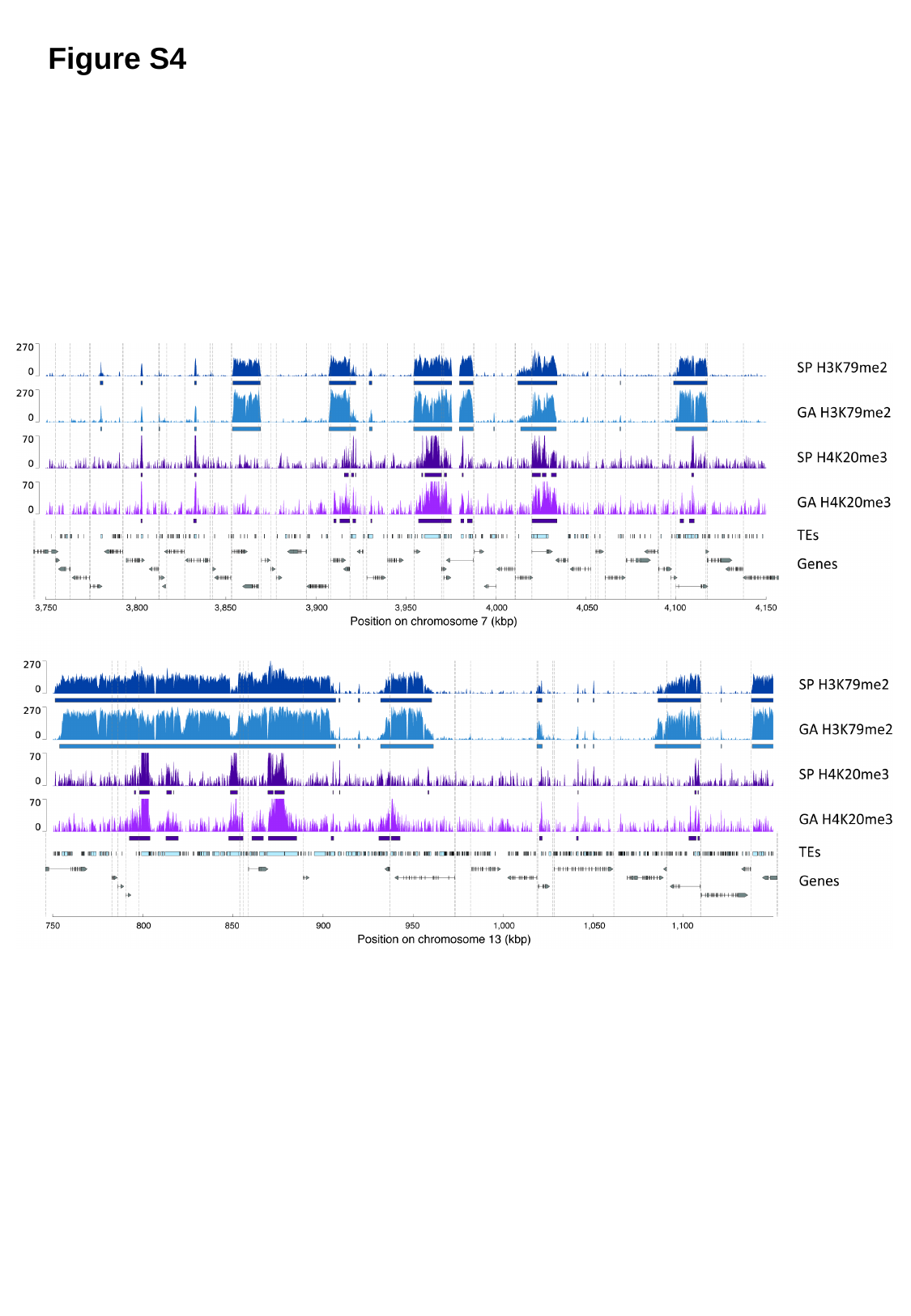

Figure S4

### Slide 3
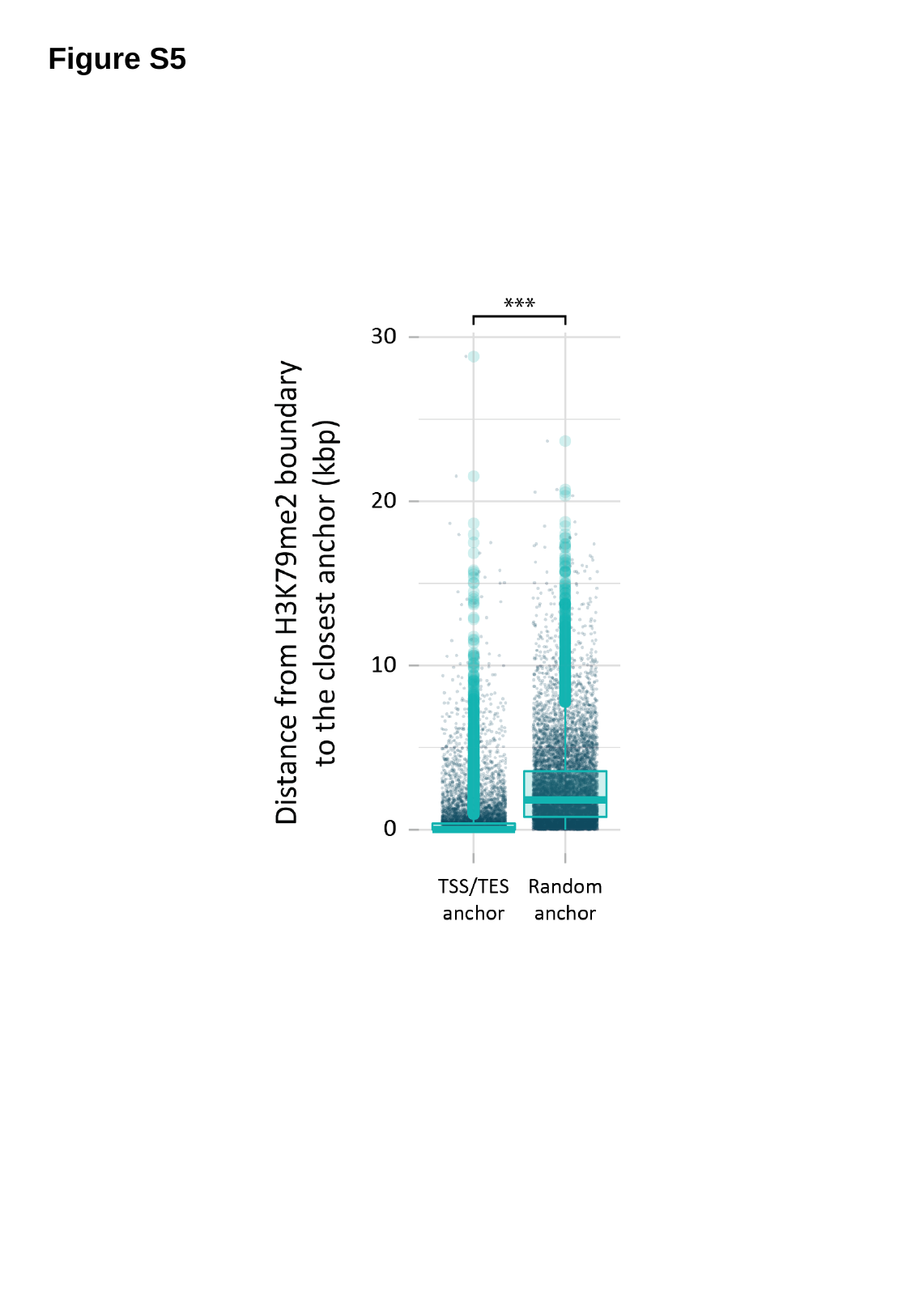

Figure S5

### Slide 4
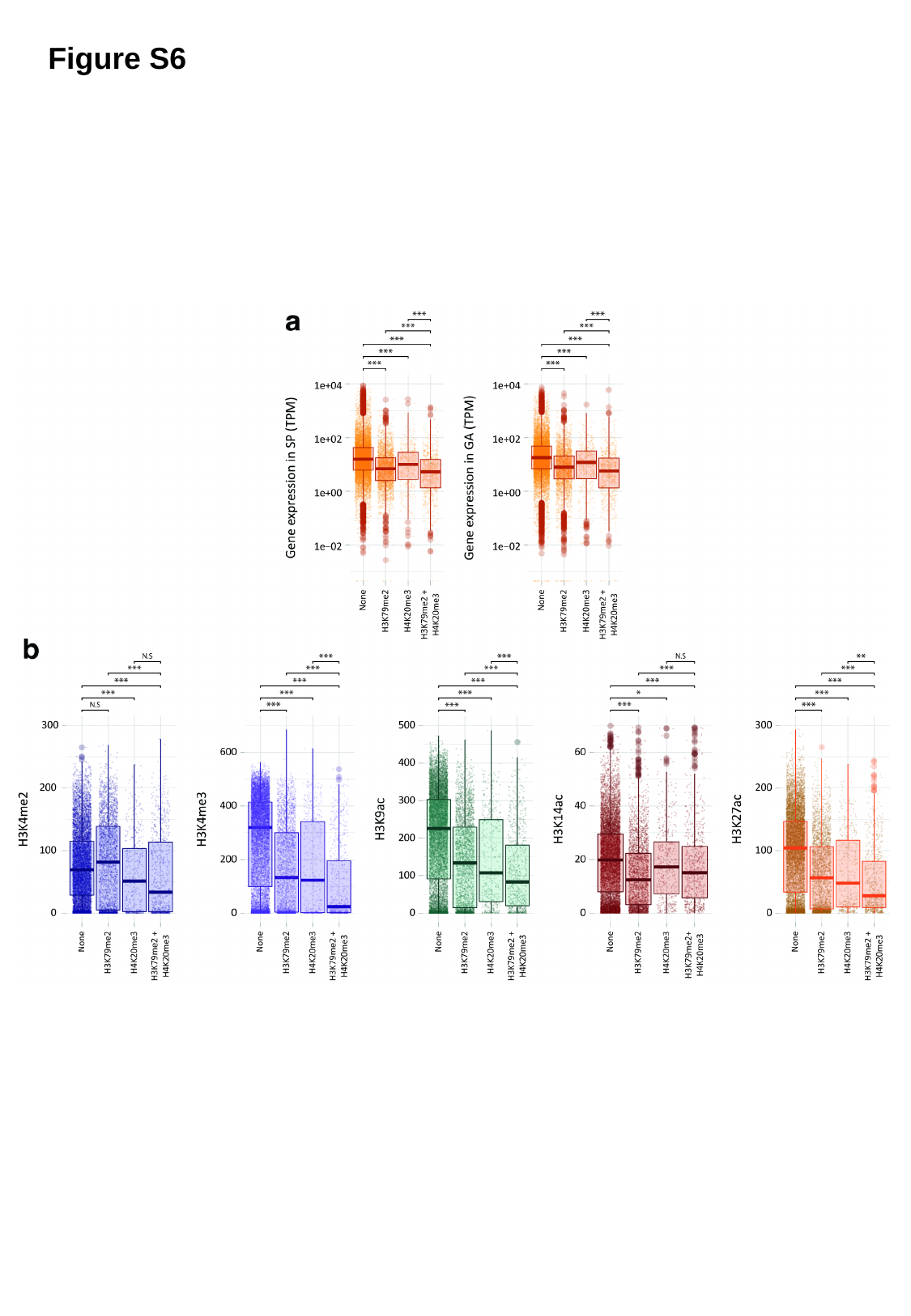

Figure S6

### Slide 5
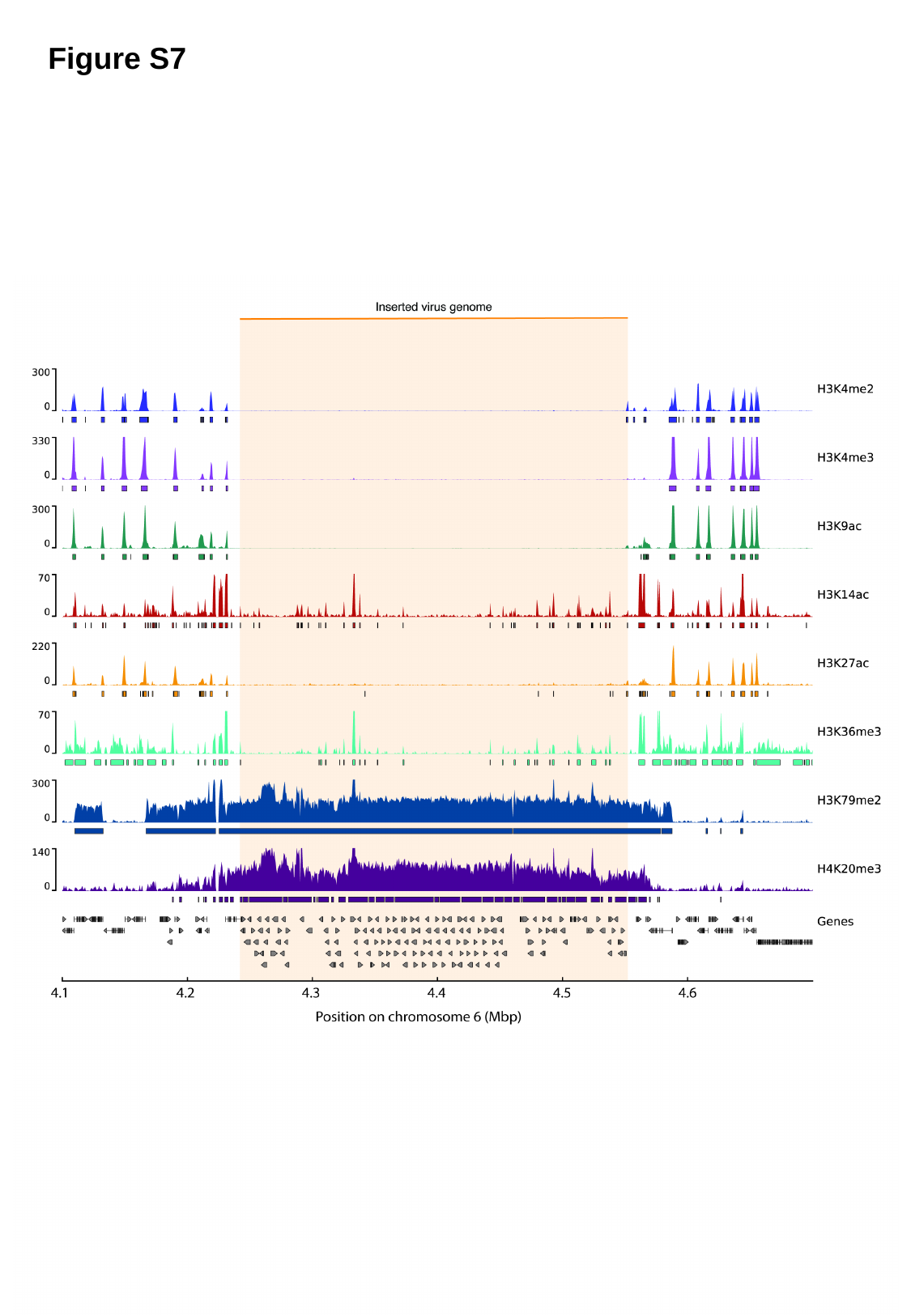

Figure S7

### Slide 6
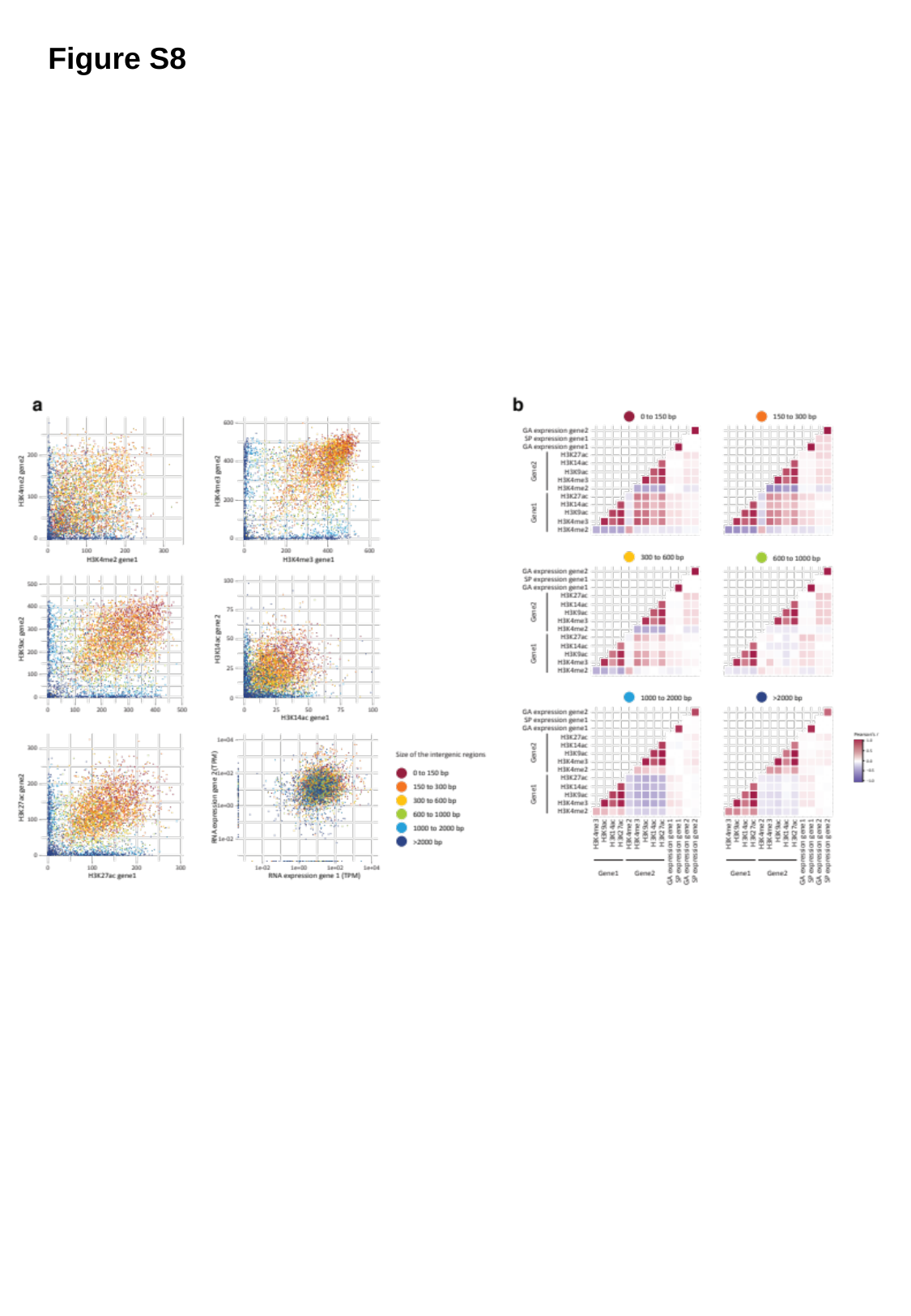

Figure S8

### Slide 7
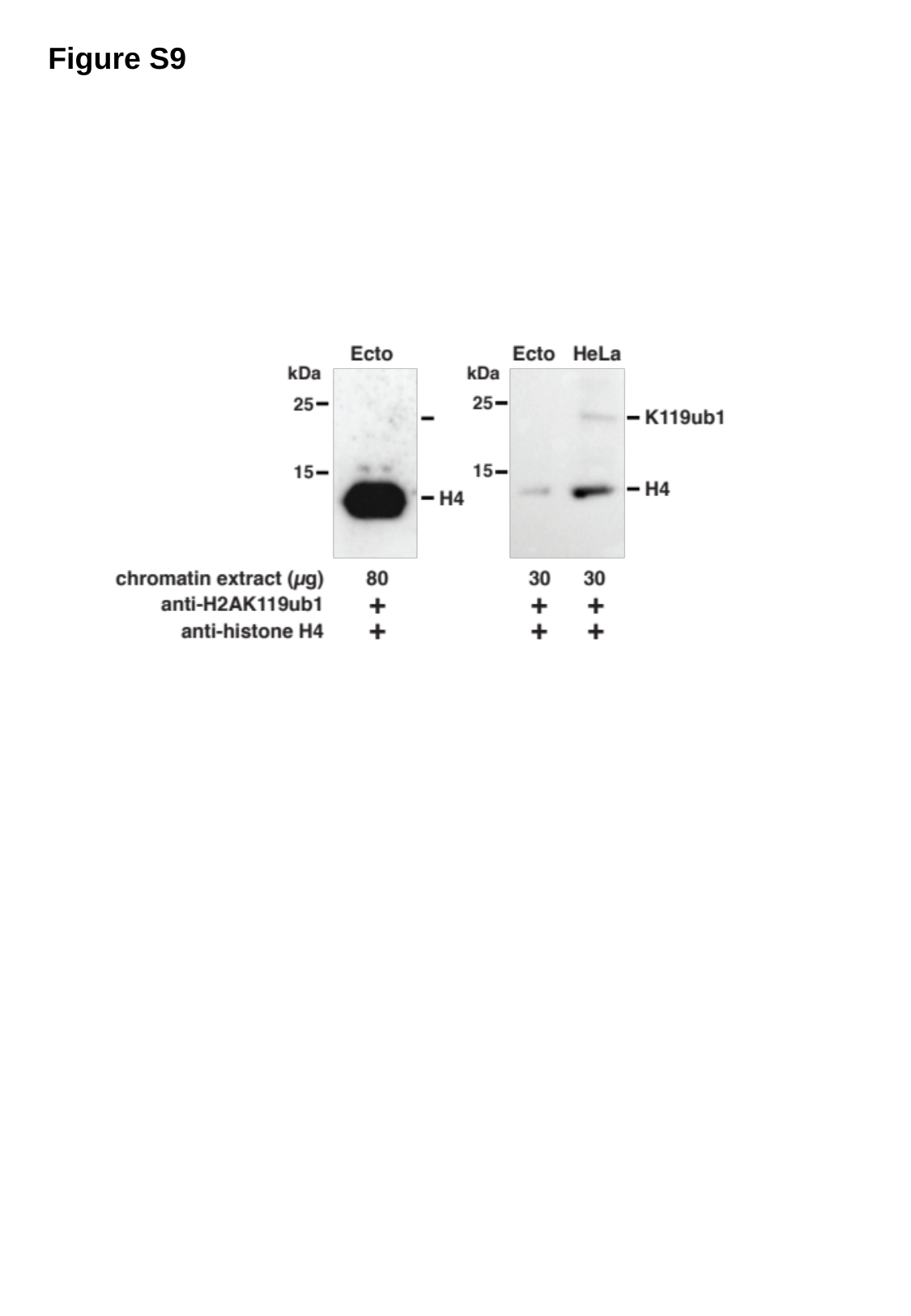

Figure S9

### Slide 8
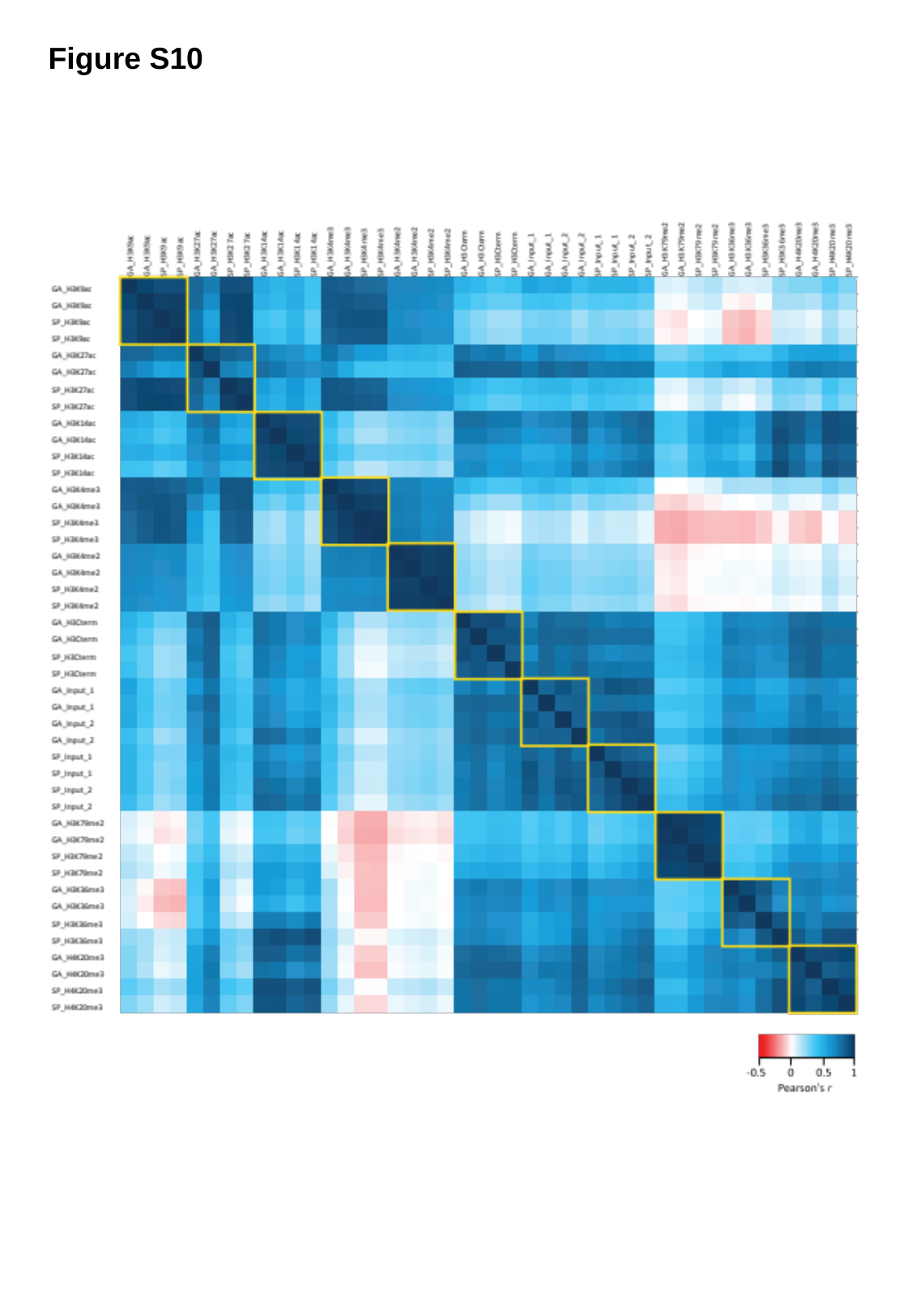

Figure S10
