## Supplementary material for "Histone modifications during the life cycle of the brown alga *Ectocarpus*": Table S2 PTM_writers_erasers

**Table S2. Putative PTM writers and erasers in *Ectocarpus* sp.** Human and *Phaeodactylum tricornutum* (diatom) orthologues are also reported. Substrate specificity and biological functions are indicated based on what is known for humans and should be only considered as putative for *Ectocarpus* enzymes. NO CAT, no known category; ψ, probable pseudogene; v, inserted viral gene.

**Table S2.1.** Histone Lysine Acetyltransferases (KAT)

| KAT | *Homo sapiens* [1] | *Ectocarpus* sp. | *Phaeodactylum tricornutum* | Substrate specificity [1] |
|---|---|---|---|---|
| *GNAT Family* | | | | |
| KAT1 | HAT1 | Ec-01_007540 | Phatr3_J54343 | H4K5 and K12 |
| KAT2A/B | GCN5 | Ec-12_003920 | Phatr3_J2957 | H3K9, K14 and K18 |
| ? | ATAC2 | No orthologues | No orthologues | H3K9, K14 and K18 |
| KAT9 | ELP3 | Ec-12_002990 | Phatr3_J50848 | H3 and maybe H4 |
| NO CAT | | Ec-01_001170, Ec-01_005970, Ec-01_005990, Ec-01_006000, Ec-01_008410, Ec-02_003420, Ec-02_006170, Ec-05_004370, Ec-07_002520, Ec-08_002320, Ec-08_005180, Ec-08_005960, Ec-15_004510, Ec-16_003580, Ec-17_000040, Ec-17_000920, Ec-17_001110, Ec-22_002740, Ec-24_004370, Ec-28_000410 | | |
| *MYST Family* | | | | |
| KAT5 | TIP60 | No orthologues | No orthologues | H4K5, K8, K12 and K16 |
| KAT6A | MOZ | No orthologues | No orthologues | H3K9 and K14 |
| KAT6B | MORF | No orthologues | No orthologues | H3K9 and K14 |
| KAT7 | HBO1 | No orthologues | No orthologues | H4K5, K8 and K12 |
| KAT8 stram_A | MOF | Ec-14_006220 | Phatr3_J51406 | H3K14 and K23, H4K16 |
| KAT8 stram_B | MOF | Ec-22_002080 | Phatr3_J3062 | |
| *p300/CBP Family* | | | | |
| KAT3A | CBP | | | Not discriminating |
| KAT3B | p300 | | | Not discriminating |
| KAT3 stram_A | | Ec-18_002450, Ec-04_001240 | Phatr3_J45703, Phatr3_J54505, Phatr3_J45764 | Not discriminating |
| KAT3 stram_B | | Ec-04_005740 | Phatr3_Jdraft292 | Not discriminating |

**Table S2.2.** Histone Lysine Methyltransferases (KMT)

| **KMT** | **Domains** | ***Ectocarpus* sp.** | ***Phaeodactylum tricornutum*** | ***Homo sapiens*** | **Substrate specificity** |
|---|---|---|---|---|---|
| ***SET domain-containing KMTs*** | | | | | |
| *MLL Family* | | | | | |
| KMT2A-D | PWWP + ZF-PHD + BRD + FY-rich + SET + Post-SET | Ec-18_000480 | Phatr3_J15937 | MLL1, MLL2, MLL3, MLL4 [2] | H3K4 [2] |
| SUV39 *Family* | | | | | |
| KMT1A-B | CHROMO + AWS + SET + Post-SET | Ec-17_000690 | No orthologues | SUV39H1, SUV39H2 [3] | H3K9 [3] |
| KMT1E-F | ZF-PHD + MBD + BRD + SET | Ec-12_006400 | No orthologues | SETDB1, SETDB2 [3] | H3K9 [3] |
| *SMYD Family* | | | | | |
| SMYD | SET + MYND | Ec-04_001520 | Phatr3_J1647, Phatr3_J43708 | SMYD1-5 [3] | H3K4, H4K20 ? [3] |
| *No human-related families* | | | | | |
| NO CAT | AWS + SET | Ec-27_005100 | No orthologues | No orthologues | Unknown |
| NO CAT | AWS + SET + Post-SET + ZF-PHD | Ec-05_003500 | Phatr3_J6093 | No orthologues | Unknown |
| NO CAT | SET + Post-SET | Ec-12_004700, Ec-14_005625, Ec-15_002050, Ec-15_002680, Ec-28_001240 | Phatr3_J21456 | No orthologues | Unknown |
| NO CAT | ZF-PHD + SET | Ec-12_006750ψ, Ec-12_006810, Ec-19_005100 | No orthologues | No orthologues | Unknown |
| NO CAT | ZF-PHD + Pre-SET + SET | Ec-14_005310 | No orthologues | No orthologues | Unknown |
| NO CAT | BRD + SET | Ec-12_006600 | No orthologues | No orthologues | Unknown |
| NO CAT | SET + BRK + ZF-CW + CHROMO + TPR | Ec-12_000580 | Phatr3_J44935 | No orthologues | Unknown |
| *KMTs with putative new functions* | | | | | |
| Kinase-containing SET | AWS + SET + Post-SET + Cyclin-like + Kinase | Ec-06_008170 | Phatr3_EG02211 | No orthologues | Unknown |
| ADD-containing SET | ZF-ADD + Pre-SET + SET + Post-SET | Ec-22_000420 | No orthologues | No orthologues | Unknown |
| *Other SET* | | | | | |
| NO CAT | SET | Ec-00_001310, Ec-01_004930, Ec-05_000850, Ec-06_002960, Ec-06_005230v, Ec-06_005320v, Ec-06_006470v, Ec-07_003020, Ec-07_004730, Ec-07_007100, Ec-08_006000, Ec-14_004620, Ec-14_004640, Ec-18_001770, Ec-19_004190, Ec-20_002460, Ec-26_000740, Ec-28_002300, Ec-14_005625 | Phatr3_J39209, Phatr3_EG01652, Phatr3_J38974, Phatr3_J50541, Phatr3_J43311, Phatr3_J43708, Phatr3_EG01005, Phatr3_J43177, Phatr3_J48703, Phatr3_J24019 | | Unknown |

| | | | | | |
|---|---|---|---|---|---|
| NO CAT | SET + Rubisco LSMT | Ec-01_001070, Ec-06_004080, Ec-11_003630, Ec-12_007560, Ec-14_001980, Ec-14_003300, Ec-20_001330 | Phatr3_J43946, Phatr3_J48815, Phatr3_J37749 | | Unknown |
| ***DOT1 domain-containing KMTs*** | | | | | |
| KMT4 | DOT1 | Ec-06_007110, Ec-12_004580, Ec-24_003550 | Phatr3_J47512, Phatr3_J44757 | DOT1 [4] | H3K79 [4] |
| KMT4_ecto | DOT1 + ZF-PHD | Ec-25_003090 | No orthologues | No orthologues | H3K79 [4] |

**Table S2.3.** Histone Arginine Methyltransferases (PRMT)

| **RMT** | ***Ectocarpus* sp.** | ***Phaeodactylum tricornutum*** | ***Homo sapiens*** |
|---|---|---|---|
| *PRMT Family* | | | |
| NO CAT | Ec-06_002620, Ec-14_006300, Ec-27_000650, Ec-27_005280 | Phatr3_J17184, Phatr3_J54710, Phatr3_J44159, Phatr3_J45331, Phatr3_EG02379 | PRMT1-9 |
| PRMT5 | Ec-10_005680 | Phatr3_J16141 | |

**Table S2.4.** Histone Deacetylases (HDAC)

| **HDAC** | ***Ectocarpus* sp.** | ***Phaeodactylum tricornutum*** | ***Homo sapiens* [5]** |
|---|---|---|---|
| *Class I* | | | |
| | Ec-15_004560, Ec-21_001720, Ec-28_001400 | Phatr3_J49800, Phatr3_J43919, Phatr3_J51026 | HDAC1-3, 8 |
| *Class II* | | | |
| | Ec-05_003890, Ec-05_006370, Ec-06_004320, Ec-11_001830, Ec-10_001980 | Phatr3_J4590, Phatr3_EG01943, Phatr3_J35869, Phatr3_J50482, Phatr3_J8891, Phatr3_J45431, Phatr3_J4423 | HDAC4,5,7,9 HDAC6, 10 |
| *Class IV* | | | |
| | Ec-14_000150, Ec-24_003590 | Phatr3_J9278, Phatr3_J4821 | HDAC11 |
| *Class III / Sirtuin Family* | | | |
| | Ec-05_003910, Ec-17_002180, Ec-19_000750, Ec-21_005430, Ec-26_003780, | Phatr3_J8827, Phatr3_J16859, Phatr3_J12305, Phatr3_J45850, Phatr3_J52135, Phatr3_J21543, Phatr3_J39523 | SIRT1-7 |

**Table S2.4.** Histone Lysine Demethylases (KDM)

| KDM | Domains | *Ectocarpus* sp. | *Phaeodactylum tricornutum* | *Homo sapiens* [6] | Substrate specificity [6] |
|---|---|---|---|---|---|
| *Lysine Specific Demethylases Family* | | | | | |
| KDM1 | Amine oxidase | Ec-10_001210, Ec-21_006430, Ec-24_001850 | Phatr3_J51708, Phatr3_EG01090, Phatr3_J44106, Phatr_J48603 | LSD1, LSD2 | H3K4me1/2, H3K9me1/2 |
| *Jumonji-C Domain-containing Family* | | | | | |
| KDM2 | F-box + JmjC | Ec-21_004490 | Phatr3_J42595 | FBXL10, FBXL11 | H3K36me1/2 |
| KDM4D | JmjN + JmjC | Ec-22_001750 | No orthologues | JMJD2 | H3K9me1/2/3 |
| KDM5A-B | JmjN +JmjC + ZF-C5HC2 + ARID + ZF-PHD | Ec-01_004230, Ec-21_005410 | Phatr3_J48747 | JARID1A-C | H3K4me2/3 |
| KDM6A | TPR + JmjC | No orthologues | No orthologues | UTX | H3K27me2/3 |
| KDM6B | JmjC | Ec-02_001410, Ec-02_001750, Ec-03_002440, Ec-07_006870, Ec-10_002180, Ec-10_005720, Ec-14_002420, Ec-14_004840, Ec-14_004970, Ec-15_003970, Ec-17_001710, Ec-21_001600, Ec-27_002280 | Phatr3_J43557, Phatr3_J48473, Phatr3_J35781, Phatr3_J42595, Phatr3_EG01348 | JMJD3 | |

### References


[1] **Yang XJ. 2016.** Histone Acetyltransferases, Key Writers of the Epigenetic Language. In *Chromatin Signaling and Diseases*, Academic Press, Elsevier.

[2] **Gu B, Lee MG. 2013.** Histone H3 lysine 4 methyltransferases and demethylases in self-renewal and differentiation of stem cells. *Cell & Bioscience* **3**: 39.

[3] **Mozzetta C, Boyarchuk E, Pontis J, Ait-Si-Ali S. 2015.** Sound of silence: the properties and functions of repressive Lys methyltransferases. *Nature Reviews Molecular Cell Biology* **16**: 499-513.

[4] **Feng Q, Wang H, Ng HH, Erdjument-Bromage H, Tempst P, Struhl K, Zhang Y. 2002.** Methylation of H3-Lysine 79 Is Mediated by a New Family of HMTases without a SET Domain. *Current Biology* **12**: 1052-1058.

[5] **Lamberti MJ, Vera RE, Rumie Vittar NB, Schneider G. 2016.** Histone Deacetylases, the Erasers of the Code. In *Chromatin Signaling and Diseases*, Academic Press, Elsevier.

[6] **García MA, Fueyo R, Martínez-Ballás MA. 2016.** Lysine Demethylases: Structure, Function, and Disfunction. In *Chromatin Signaling and Diseases*, Academic Press, Elsevier.
