## Supplementary material for "Histone modifications during the life cycle of the brown alga *Ectocarpus*": Table S3 PTM_comparison

**Table S3. Presence or absence of histone post-translational modifications in seven species from diverse eukaryotic supergroups.** Probable equivalent PTMs at slightly different positions are also indicated. Not detected, not detected by either mass spectrometry or immunoblot; *, PTM only detected by immunoblot; †, the mark H3K27me3 was also weakly detected by immunoblot but further analysis indicated that it is not present in *Ectocarpus* (see text for details); No equivalent, the species in question has no amino acid residue equivalent to the *Ectocarpus* residue; Ac, acetylation; Me1, monomethylation; Me2, dimethylation; Me3, trimethylation; Pr, propylation; Ub, ubiquitination; -, not determined; Nt, N-alpha terminal.

| **Histone residue** | ***Ectocarpus* sp.** | ***Phaeodactylum tricornutum* [1]** | ***Thalassiosira pseudonana* [2]** | ***Tetrahymena thermophila* [3,4]** | ***Arabidopsis thaliana* [5–7]** | ***Homo sapiens* [8–12]** | ***Saccharomyces cerevisiae* [4,10,11,13]** |
|---|---|---|---|---|---|---|---|
| *Histone H2A canonical* | | | | | | | |
| S1 | Ac(Nt) | _ | _ | _ | _ | Ac(Nt) | Ac(Nt) |
| K3 | Ac | Ac | Not detected | No K3 equivalent | Not detected | No K3 equivalent | K4 Ac |
| K5 | Ac | Ac, K7Ac | Not detected | Ac, K8 Ac, K10 Ac, K12 Ac | Ac | Ac, K9Ac | K7 Ac |
| *Histone H2A.Z* | | | | | | | |
| S1 | Ac(Nt) | _ | _ | _ | _ | Not detected | _ |
| K3 | Ac | Ac | _ | _ | _ | K4 Ac | _ |
| K6 | Ac | Ac | _ | _ | _ | K7 Ac | _ |
| K9 | Ac | Ac | _ | _ | _ | No K9 equivalent | _ |
| K12 | Ac | Ac | _ | _ | _ | K11 Ac | _ |
| K15 | Ac | Ac | _ | _ | _ | Not detected | _ |
| K20 | Ac | Not detected | _ | _ | _ | K13 Ac | _ |
| R38 | Me1 | Not detected | _ | _ | _ | Not detected (R31) | _ |
| *Histone H2B* | | | | | | | |
| K2 | Ac | Ac | Ac | K3 not detected, Me3 | No K2 equivalent | No K2 equivalent | K3 Ac |
| K6 | Ac | Ac | Ac | K4 Ac | Ac | K5 Ac | Ac |
| K7 | Ac | K6 Ac | K6 Ac | K5 not detected | K6 Ac | K5 Ac | Ac |
| K10 | Ac | Ac | K11 Ac | No K10 equivalent | K11 Ac | K11 Ac, K12 Ac | K11 Ac |
| K13 | Ac | Ac | K14 Ac | K12 not detected | K12 not detected | K15 Ac | K16 Ac |
| K14 | Ac | Ac | K15 Ac | K13 not detected | K12 not detected | K16 Ac | Not detected |
| K34 | Not detected | Ac | No K34 equivalent | No K34 equivalent | K32 Ac, K27 Ac | Not detected | K21 Ac, K22 Ac |
| K37 | Not detected | K38 not detected | K38 Ac | K38 Not detected | K32 Ac | No K37 equivalent | Not detected |

| | | | | | | | |
|---|---|---|---|---|---|---|---|
| K47 | Not detected | K48 Ac | K52 Ac, Me2 | No K47 equivalent | K50 not detected | K47 Me1 | K46 not detected |
| K107 | Not detected | Ac | K103 Ac | No K107 equivalent | K105 not detected | K108 Me1 | K108 Ac |
| K111 | Ub | Ub | Ub not detected, Ac | Ub not detected, Ac | K143 Ub | K120 Ub | K123 Ub |
| *Histone H3* | | | | | | | |
| K4 | Me2*, Me3* | Me1, Me2, Me3 | Not detected | Not detected | Me1, Me2, Me3 | Me1, Me3 | Me1, Me2, Me3, Ac |
| K9 | Ac | Ac | Ac | Ac | Ac | Ac | Ac |
| K9 | Me1*, Me2*, Me3* | Me2*, Me3* | Not detected | Not detected | Me1, Me2*, Me3* | Me1, Me2, Me3 | Me2, Me3 |
| K14 | Ac | Ac | Ac | Ac | Ac | Ac, Me1 | Ac, Me2 |
| K18 | Ac | Ac | Ac | Ac | Ac | Ac, Me1 | Ac, Me1 |
| K23 | Ac | Ac | Ac | Ac, Me1 | Ac | Ac, Me1 | Ac, Me1 |
| K27 | Ac | Ac | Ac | Ac | Ac | Ac | Ac |
| K27 | Me2† | Me1, Me2, Me3 | Me2 | Me1, Me2, Me3 | Me1, Me2, Me3 | Me1, Me2, Me3 | Me1, Me2, Me3 |
| K36 | Not detected | Ac | Ac | Ac | Ac | Not detected | Ac |
| K36 | Me1, Me2, Me3 | Me1, Me2, Me3 | Me1, Me2, Me3 | Me1 | Me1, Me2, Me3 | Me1, Me2, Me3 | Me1, Me2, Me3 |
| K56 | Not detected | Ac | Not detected | Ac | Not detected | Ac, Me1, Me3 | Ac |
| K79 | Not detected | Ac | Ac | Not detected | Not detected | Ac | Not detected |
| K79 | Me1, Me2 | Me1, Me2 | Me1, Me2, Me3 | Me1 | Not detected | Me1, Me2, Me3 | Me1, Me2, Me3 |
| K122 | Not detected | Ac | Ac | Not detected | Not detected | Ac, Me1 | Not detected |
| *Histone H4* | | | | | | | |
| S1 | Ac(Nt) | _ | _ | _ | _ | Ac(Nt) | Ac(Nt) |
| K5 | Ac | Ac | Ac | K4 Ac | Ac | Ac, Me1 | Ac |
| K5 | Pr | _ | _ | _ | _ | Pr | _ |
| K8 | Ac | Ac | Ac | K7 Ac | Ac | Ac | Ac, Me1 |
| K12 | Ac | Ac | Ac | K11 Ac | Ac | Ac | Ac, Me1 |
| K16 | Ac | Ac | Ac | K15 Ac | Ac | Ac | Ac |
| K20 | Me1, Me2, Me3* | Ac, No methylation detected | Ac, No methylation detected | Me1 | Ac, Me3 | Me1, Me2, Me3 | Me1, Me2 |
| K31 | Not detected | Ac | Not detected | Not detected | Not detected | Me2 | Ac |
| R55 | Not detected | Not detected | Not detected | Not detected | Not detected | Me1, Me2 | Not detected |
| K59 | Not detected | Ac | Ac | Not detected | Not detected | Not detected | Not detected |
| K59 | Me1, Me2 | Me1 | Me1 | Not detected | Not detected | Me1, Me2 | Me1 |

| | | | | | | | |
|---|---|---|---|---|---|---|---|
| K79 | Me2, Me3 | Me1, Me2, Me3 | Me1, Me2, Me3 | R77 Me1 | Not detected | me2, K77 Me1 | K77 Me1 |
| K91 | Not detected | Not detected | Ac | Not detected | Not detected | Ac | Not detected |

### References


1. Veluchamy A, Rastogi A, Lin X, Lombard B, Murik O, Thomas Y, et al. An integrative analysis of post-translational histone modifications in the marine diatom *Phaeodactylum tricornutum*. Genome Biol. 2015;16:102.

2. Rastogi A, Lin X, Lombard B, Loew D, Tirichine L. Probing the evolutionary history of epigenetic mechanisms: what can we learn from marine diatoms. AIMS Genet. 2.

3. Zhang C, Gao S, Molascon AJ, Wang Z, Gorovsky MA, Liu Y, et al. Bioinformatic and proteomic analysis of bulk histones reveals PTM crosstalk and chromatin features. J Proteome Res. 2014;13:3330–7.

4. Morris SA, Rao B, Garcia BA, Hake SB, Diaz RL, Shabanowitz J, et al. Identification of histone H3 lysine 36 acetylation as a highly conserved histone modification. J Biol Chem. 2007;282:7632–40.

5. Zhang K, Sridhar VV, Zhu J, Kapoor A, Zhu J-K. Distinctive core histone post-translational modification patterns in *Arabidopsis thaliana*. PLoS One. 2007;2:e1210.

6. Charron J-BF, He H, Elling AA, Deng XW. Dynamic landscapes of four histone modifications during deetiolation in *Arabidopsis*. Plant Cell. 2009;21:3732–48.

7. Mahrez W, Arellano MST, Moreno-Romero J, Nakamura M, Shu H, Nanni P, et al. H3K36ac Is an Evolutionary Conserved Plant Histone Modification That Marks Active Genes. Plant Physiol. 2016;170:1566–77.

8. Beck HC, Nielsen EC, Matthiesen R, Jensen LH, Sehested M, Finn P, et al. Quantitative Proteomic Analysis of Post-translational Modifications of Human Histones. Mol Cell Proteomics. American Society for Biochemistry and Molecular Biology; 2006;5:1314–25.

9. Tan M, Luo H, Lee S, Jin F, Yang JS, Montellier E, et al. Identification of 67 histone marks and histone lysine crotonylation as a new type of histone modification. Cell. 2011;146:1016–28.

10. Zhao Y, Garcia BA. Comprehensive Catalog of Currently Documented Histone Modifications. Cold Spring Harb Perspect Biol. 2015;7:a025064.

11. Hole K, Van Damme P, Dalva M, Aksnes H, Glomnes N, Varhaug JE, et al. The human N-alpha-acetyltransferase 40 (hNaa40p/hNatD) is conserved from yeast and N-terminally acetylates histones H2A and H4. PloS One. 2011;6:e24713.

12. Huang H, Sabari BR, Garcia BA, Allis CD, Zhao Y. SnapShot: Histone Modifications. Cell. 2014;159:458-458.e1.

13. Valero ML, Sendra R, Pamblanco M. Tandem affinity purification of histones, coupled to mass spectrometry, identifies associated proteins and new sites of post-translational modification in Saccharomyces cerevisiae. J Proteomics. 2016;136:183–92.
